# Tissue-specific actions of Pax6 on proliferation-differentiation balance in the developing forebrain are Foxg1-dependent

**DOI:** 10.1101/374074

**Authors:** Idoia Quintana-Urzainqui, Zrinko Kozić, Soham Mitra, Tian Tian, Martine Manuel, John O. Mason, David J. Price

**Affiliations:** Simons Initiative for the Developing Brain, Hugh Robson Building, George Square, Edinburgh EH8 9XD, UK

**Keywords:** transcription factors, forebrain development, Pax6, Foxg1, cerebral cortex, thalamus, prethalamus, diencephalon, telencephalon, proliferation, differentiation

## Abstract

Differences in the growth and maturation of diverse forebrain tissues depends on region-specific transcriptional regulation. Individual transcription factors act simultaneously in multiple regions that develop very differently, raising questions about the extent to which their actions vary regionally. We found that the transcription factor Pax6 affects the transcriptomes and the balance between proliferation and differentiation in opposite directions in murine diencephalon versus cortex. We tested several possible mechanisms to explain Pax6’s tissue-specific actions and found that the presence of the transcription factor Foxg1 in cortex but not diencephalon was most influential. We found that Foxg1 is responsible for many of the differences in cell cycle gene expression between diencephalon and cortex. In cortex lacking Foxg1, Pax6’s action on the balance of proliferation versus differentiation became diencephalon-like. Our findings reveal a mechanism for generating regional forebrain diversity in which the actions of one transcription factor completely reverse the actions of another.

## Introduction

The mechanisms that create the brain’s enormous interregional diversity of structure and function remain poorly understood. Early in embryogenesis, the anterior neural plate is patterned by the regional expression of transcription factors whose actions are essential for each part to acquire its correct size and cellular composition. These transcription factors are sometimes referred to as “master regulators” or “selectors” to reflect their powerful ability to control the hierarchies of gene expression that specify region-specific growth and identity (Allan and Thor, 2015). Individual transcription factors do not instruct the development of unique regions but are expressed simultaneously in multiple regions whose morphologies and functions develop very differently. This raises questions about the degree to which the actions and the mechanisms of action of individual transcription factors vary between different brain regions. Does a transcription factor regulate a particular process, such as cell proliferation, similarly in all regions? To what extent and how are its actions modified by the context in which it is expressed?

The transcription factor Pax6 is expressed simultaneously by large populations of progenitors in both of the forebrain’s major components, the telencephalon and the diencephalon (Stoykova and Gruss, 1994). During this time, the telencephalon expands much more than the diencephalon, which it engulfs as it forms the cerebral cortex dorsally and the basal ganglia ventrally. The diencephalon forms the thalamus (Th) and prethalamus (PTh), which process and transmit signals to and from the overlying cortex. Pax6 regulates the proliferation of cortical cells by limiting the rate at which progenitors progress through the cell cycle. Pax6 deletion promotes a higher rate of proliferation of cortical radial glial progenitors (RGPs) whereas the opposite occurs if Pax6 is overexpressed within a physiological range (Georgala et al., 2011a, 2011b; Manuel and Price, 2005; Mi et al., 2013). Previous studies have shown that Pax6 is required for normal thalamic and prethalamic development (Stoykova et al., 1996; Warren and Price, 1997), but whether it controls thalamic and prethalamic progenitors in the same way it regulates cortical RGPs is unknown.

We began by addressing this question. We compared the effects of acute Pax6 deletion on the transcriptomes of cells from embryonic cortex, Th and PTh and found that the expression levels of genes associated with cell proliferation and differentiation were altered in opposite directions in cortex and diencephalon. We went on to show that this corresponded with an opposite effect of Pax6 deletion on the balance between proliferation and differentiation in these two regions. We next explored mechanisms that might account for these tissue-specific differences, including the possibility that the actions of Pax6 are affected by the presence or absence of another high-level transcription factor, Foxg1. Foxg1 regulates the cell cycles of cortical progenitors, is expressed by telencephalic cells including RGPs but is not expressed by diencephalic cells (Fasano et al., 2009; Kumamoto and Hanashima, 2017; Mariani et al., 2016; Martynoga et al., 2005; Xuan et al., 1995). We first showed that Foxg1 deletion from cortical cells makes their profile of cell cycle gene expression more diencephalon-like. We went on to test how *Foxg1*-null cortical cells respond to Pax6 deletion. We found this caused cell-autonomous changes that were opposite to those in Foxg1-expressing cortex and similar to those in diencephalic tissues, indicating that the region-specific actions of Pax6 on the balance of proliferation and differentiation in developing forebrain are Foxg1-dependent.

## Results

### Major inter-regional differences of identity remain after Pax6 loss from forebrain

We started by using RNA-seq to study the effects of tamoxifen-induced *Pax6* deletion on gene expression in cortex, Th and PTh at embryonic day 13.5 (E13.5). Administration of tamoxifen at E9.5 to *Pax6^fl/fl^* embryos ubiquitously expressing Cre recombinase from a *CAG^CreER^* allele caused Pax6 loss from E11.5 onwards (Figure 1A-D). These embryos are referred to here as *CAG^CreER^ Pax6* cKOs (conditional knockouts) and they are compared to *CAG^CreER^ Pax6^fll+^* littermate controls, which continue to express Pax6 in a normal pattern (Figure 1A,C). Heterozygosity for *Pax6* does not detectably affect forebrain Pax6 protein levels nor the proliferation of Pax6 expressing cells (Figure 1-figure supplement 1A; Mi et al., 2013). Accurate and consistent dissection of the Th, PTh and the anterior half of the cortex (ACtx) (where the cortical level of Pax6 is highest) was guided by the *DTy54* transgene, which expresses green fluorescent protein (GFP) under the control of all known *Pax6* regulatory elements (Tyas et al., 2006a). This transgene faithfully reports levels of *Pax6* gene expression in cells in which the endogenous *Pax6* locus can be either normal or null. PTh and ACtx are distinguished by high levels of *Pax6/*GFP expression; PTh has a sharp posterior boundary with Th, which expresses *Pax6/*GFP at lower levels (Figure 1-figure supplement 1B-E’).

**Figure 1.**
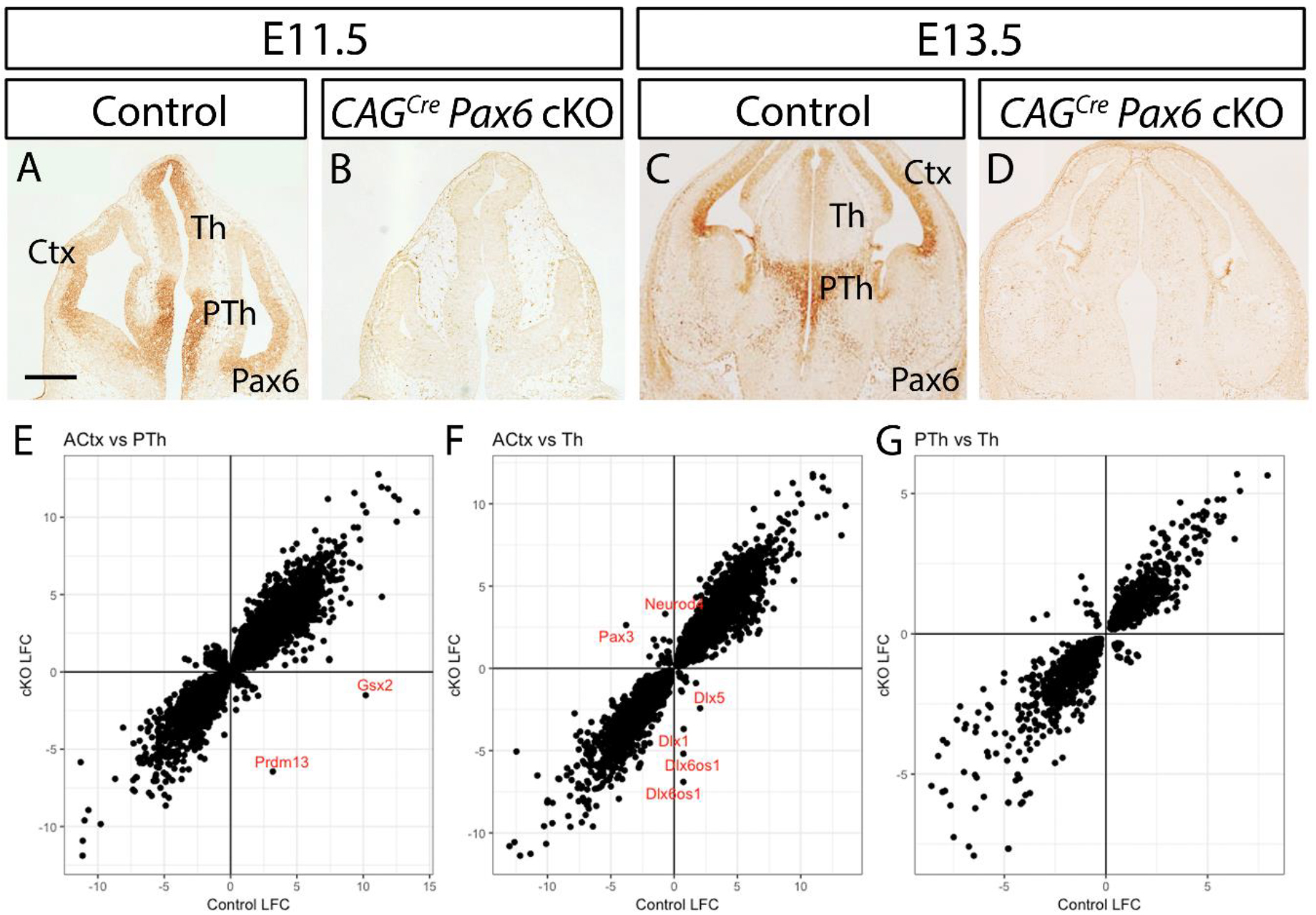
Effects of Pax6 deletion on transcriptomic differences between embryonic anterior cortex (ACtx), thalamus (Th) and prethalamus (PTh). (A-D) Pax6 immunohistochemistry at E11.5 and E13.5 showing *CAG^CreER^*-induced loss of Pax6 throughout the forebrain following tamoxifen administration at E9.5. Ctx: cortex. Scale bar: 0.25mm. (E-G) Log_2_ fold change (LFC) of gene expression between (E) ACtx and PTh, (F) ACtx and Th and (G) PTh and Th in controls and *CAG^CreER^ Pax6* cKOs. Positive values indicate enrichment in PTh in (E) and in Th in (F,G). Negative values indicate enrichment in ACtx in (E,F) and in PTh in (G). Selected genes are labelled in red. Linked to Figure 1–figure supplement 1, Figure 1-figure supplement 2 and Supplementary Files 1,2 and 3.

Prior to RNA-seq, we quality-controlled the accuracy and consistency of the separation of PTh and Th using quantitative real-time PCR (qRT-PCR) to measure the levels of expression of *Dlx2*, which is highly expressed in PTh but not in Th, and *Neurog2*, which is highly expressed in Th and in only a small proportion of PTh cells (Figure 2-figure supplement 1A,B). None of the PTh samples included in the analysis contained mRNA for *Wnt8b*, confirming that they were not contaminated by adjacent eminentia thalami, which is *Wnt8b*-rich (Adutwum-Ofosu et al., 2016). At least three quality-controlled replicate samples representing each tissue and genotype from independent litters were analysed by RNA-seq. After sequencing, we re-confirmed the accuracy of our diencephalic dissections by extracting the expression values of several reference genes with well-characterized differential expression between the PTh and Th. We found low levels of prethalamic markers (*Dlx2*, *Gsx2* and *Ascl1*) in both control and cKO thalamic samples (red arrows in Figure 2-figure supplement 1C,E,G) and low levels of thalamic markers (*Neurog2*, *Gbx2* and *Dbx1*) in both control and cKO prethalamic samples (red arrows in Figure 2-figure supplement 1D,F,H). The raw data from the RNA-seq is publically available from the European Nucleotide Archive (ENA) (www.ebi.ac.uk/ena; Project 2015054, ENA accessions PRJEB9747, ERP010887). Principal component analyses (PCA) showed that individual samples separated according to genotype along the axes of greatest variation in all three tissues (Figure 2-figure supplement 2).

We first examined the differences between the transcriptomes of ACtx, Th and PTh in control embryos. Supplementary File 1 lists all genes showing significant (adjusted p<0.05) enrichment in each control tissue over its level in each of the other two control tissues. There were about 3,000 significant differences between Th and PTh and about 10-11,000 between ACtx and Th or PTh. The expression patterns of genes showing the greatest inter-regional differential expression are shown in Figure 1-figure supplement 2. Many of these genes encode transcription factors or other molecules known to be involved in the regulation of developmental processes. This analysis illustrates the enormous divergence in molecular identities and hence the context within which Pax6 operates in these three regions.

We then repeated this analysis on data from *CAG^CreER^ Pax6* cKOs to compare the differences between the transcriptomes of ACtx, Th and PTh when Pax6 was deleted. The numbers of inter-regional differences increased to over 4,000 between Th and PTh and to over 12,000 between ACtx and either Th or PTh (Supplementary File 2). We then paired the regions - ACtx with Th, ACtx with PTh, Th with PTh - and calculated differential expression (in the form of log_2_ fold changes, LFCs) between the members of each pair in controls and in cKOs. For each pair, we then plotted control LFCs and cKO LFCs against each other, including all genes significantly differentially regulated by Pax6 loss in both members of each pair (Figure 1E-G). The graphs showed that the vast majority of genes retained the direction of their inter-regional differential expression in cKOs (i.e. if they were higher in one region in controls, the same was true in cKOs). Exceptions included *Gsx2*, which is known from previous work to have its normally strong expression in PTh extinguished by Pax6 loss (Caballero et al.), and Dlx family members, which are upregulated in cKO ACtx (Figure 1E,F; all exceptions are listed in Supplementary File 3). We conclude that Pax6 is not required to maintain the vast majority of fundamental differences in molecular identity between these forebrain regions.

### The sets of genes regulated by Pax6 vary between forebrain regions

We next studied gene expression changes between controls and *CAG^CreER^ Pax6* cKOs within each individual tissue. Supplementary File 4 lists, for each tissue, all genes showing significant differential expression between controls and cKOs (adjusted p<0.05). Figure 2A-C plots, for each tissue, each gene’s LFC in expression over its average expression (those with adjusted p<0.05 in red) and Figure 2D summarizes the numbers of significantly up- and down-regulated genes in cKOs. To explore the lists in Supplementary File 4 interactively, visit *Differential Expression* at https://pricegroup.sbms.mvm.ed.ac.uk/Pax6_diencephalon/. PTh showed the greatest number of differentially expressed genes, probably because *Pax6* is expressed not only by all progenitors but also by many postmitotic neurons in PTh, whereas it is expressed only by progenitors in cortex and Th. This likelihood is supported by the observation that genes showing the largest differential expression values in PTh include many encoding ion channels and receptors associated with postmitotic neurons, whereas these genes showed little or no significant differential expression in ACtx or Th (Supplementary File 4). Quite similar numbers of genes showed significant differential expression in ACtx and Th (Figure 2D). The numbers of significantly upregulated genes were ^~^40-60% higher than the numbers of downregulated genes in both Th and Pth, whereas this ratio was reversed in ACtx.

**Figure 2.**
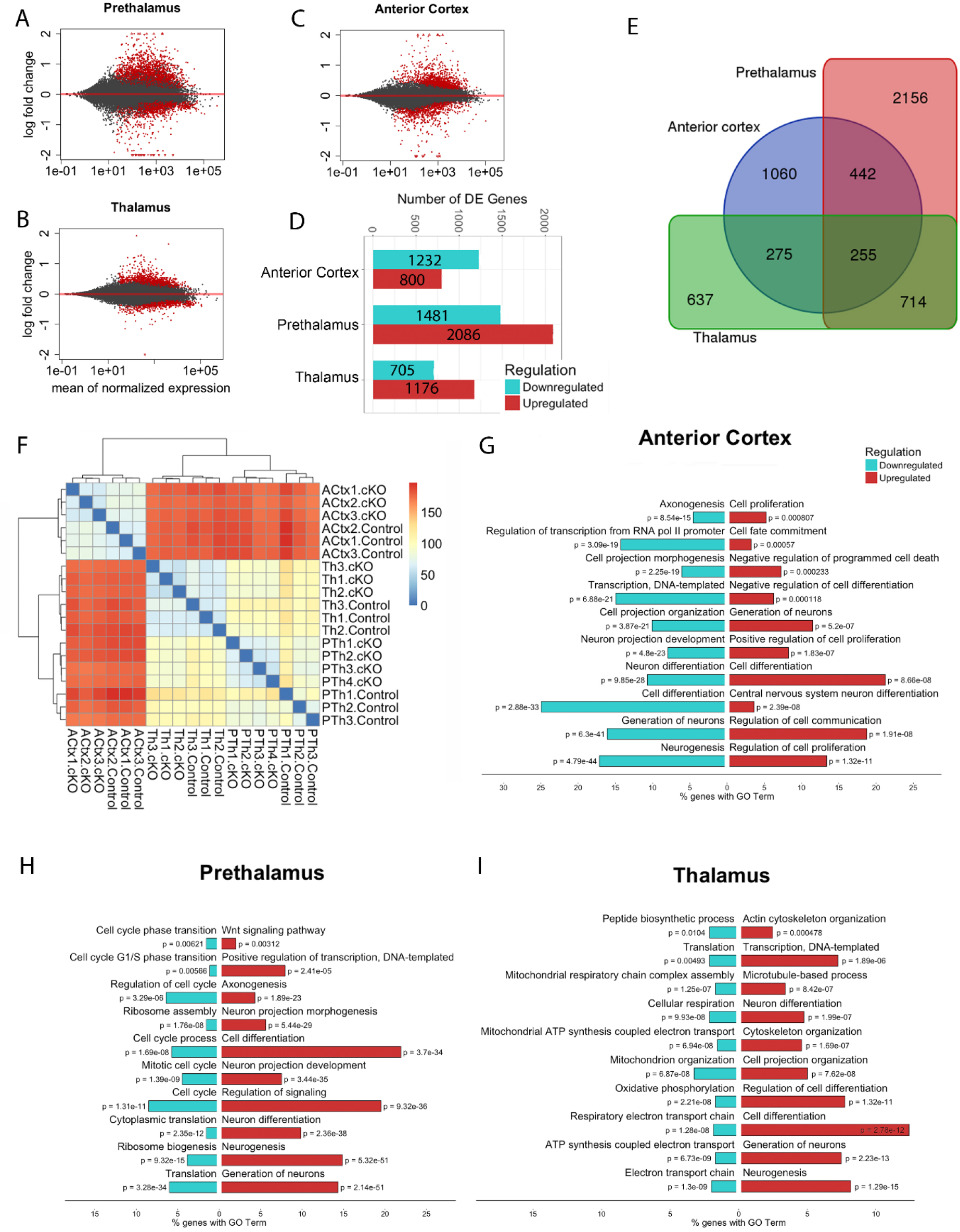
The effects of Pax6 deletion on the transcriptomes of prethalamic, thalamic and anterior cortical cells. (A-C) MA plots of log_2_ fold changes in the expression of each gene against its average expression level; red dots indicate statistically significant changes between genotypes (adjusted p values <0.05). (D) An overview of the numbers of significantly differentially expressed (DE) genes resulting from Pax6 deletion in each region. (E) Venn diagram showing the numbers of significantly DE genes in each region and common to multiple regions. (F) Heatmap representing the results of hierarchical clustering of RNAseq data from each sample from control and *CAG^CreER^ Pax6* cKO. (G-I) The ten most highly enriched, non-redundant gene ontology (GO) terms with obvious relevance to developmental processes for upregulated and downregulated genes in each region. Linked to Figure 2–figure supplement 1, Figure2-figure supplement 2 and Supplementary Files 4 and 5.

A surprisingly small number of genes (255 out of 5539) showed significant differential expression across all three tissues (referred to here as “commonly deregulated genes”) (Figure 2E). Forty-five of these genes were upregulated and only 12 were downregulated in all three tissues (listed in Supplementary File 5). Many of these 57 genes encode molecules implicated in intercellular interactions such as cell adhesion, cell-cell communication and axon guidance. For example, proteins involved in Wnt signalling were strongly represented. *Wnt3a*, *Wnt5a*, *Lrg4*, *Wif1* and *Wls* were commonly upregulated and *Dkk3* was commonly downregulated. Among the commonly downregulated genes were two related to retinoic acid signalling: *Rlbp1* and *Ripply3*. To see the expression levels of any regulated gene in our dataset visit *Gene Expression Plots* at https://pricegroup.sbms.mvm.ed.ac.uk/Pax6_diencephalon/. Interestingly, 8 of the 12 genes that are commonly downregulated have been reported to be directly regulated by Pax6 (*Dkk3*, *Rlbp1*) and/ or show a peak in Pax6 chromatin immunoprecipitation (ChIP)-seq (*Bmpr1b*, *Cldn12*, *Dkk3*, *Mlc1*, *Nr2e1*, *Rypply3*, *Rlbp1* and *Sema3a*; references in Supplementary File 5). The scarcity of commonly deregulated genes and the fact that the direction of change of 198/255 of them (i.e. whether they were up- or down-regulated) varied between tissues suggests that Pax6 deletion affects aspects of cortical, thalamic and prethalamic development in substantially different ways.

### Pax6 loss affects gene expression profiles oppositely in cortex and diencephalon

To discover global similarities and differences in the inter-regional effects of Pax6 on gene expression, we carried out distance-based hierarchical clustering on our RNA-seq data. This identified two major clusters representing samples from cortex and diencephalon (Figure 2F), suggesting that the reactions of the two diencephalic tissues, Th and PTh, to Pax6 loss might be more similar to each other than to that of the cortex.

To predict how Pax6 removal might affect biological processes in ACtx, Th and PTh and to look for possible similarities and differences in how these regions respond to the deletion, we first carried out gene ontology (GO) term enrichment analysis on all genes showing significant differential expression in each tissue. Figure 2G-I shows the ten most highly enriched, non-redundant GO terms with obvious relevance to developmental processes for upregulated and downregulated genes in each tissue. Regarding genes upregulated in cortex, 3/10 terms contained “cell proliferation” with one being “positive regulation of cell proliferation” and 3/10 terms contained “differentiation” with one being “negative regulation of cell differentiation”. Regarding genes downregulated in cortex, no terms related to proliferation but 2/10 terms contained “differentiation” and a further 4/10 terms related to processes that occur in differentiating cells, such as “axonogenesis” or “projection development”. By contrast, in diencephalic tissues terms including “differentiation”, “axonogenesis” or “projection development” were associated only with upregulated genes (5/10 in PTh and 4/10 in Th) whereas terms related to proliferation were associated only with downregulated genes (6/10 terms contained “cell cycle” in PTh). Overall, this analysis suggests that the effects of Pax6 loss on at least some developmental processes are likely to be opposite in cortex and diencephalon. Pax6 loss might promote proliferation in the former but promote differentiation in the latter.

We tested this idea further by hierarchically clustering all genes showing significant differential expression in at least one tissue according to the direction and magnitude of their LFC across all three tissues (Figure 3A). Most genes showed LFCs in the same direction in PTh and Th, but in an opposite direction in ACtx. The dendrogram was cut to produce 14 clusters suitable for GO term analysis. Figure 3A lists highly enriched, representative functional terms alongside each cluster (Supplementary File 6 contains a complete list of functional terms associated with each cluster). In Clusters 1 and 4, most of the genes showed positive LFCs in ACtx but negative LFCs in Th and PTh. These clusters were strongly associated with the GO term “cell cycle”. Cluster 4 contains 21 cell cycle-related terms with significant enrichment; cell cycle terms were exclusively enriched in these two clusters and not in others. Trends were opposite in clusters 6-8, with most genes showing negative LFCs in ACtx but positive LFCs in Th and PTh. These clusters were strongly associated with GO terms containing or related to “differentiation”. Many genes in clusters 5, 6 and 8, which are strongly associated with terms related to differentiation, showed the greatest LFCs in PTh, in agreement with the continued function for Pax6 in postmitotic neurons in this region. Interestingly, cluster 8 also showed enrichment for “Wnt signalling pathway”, indicating that this pathway might be upregulated specifically in the diencephalon.

**Figure 3.**
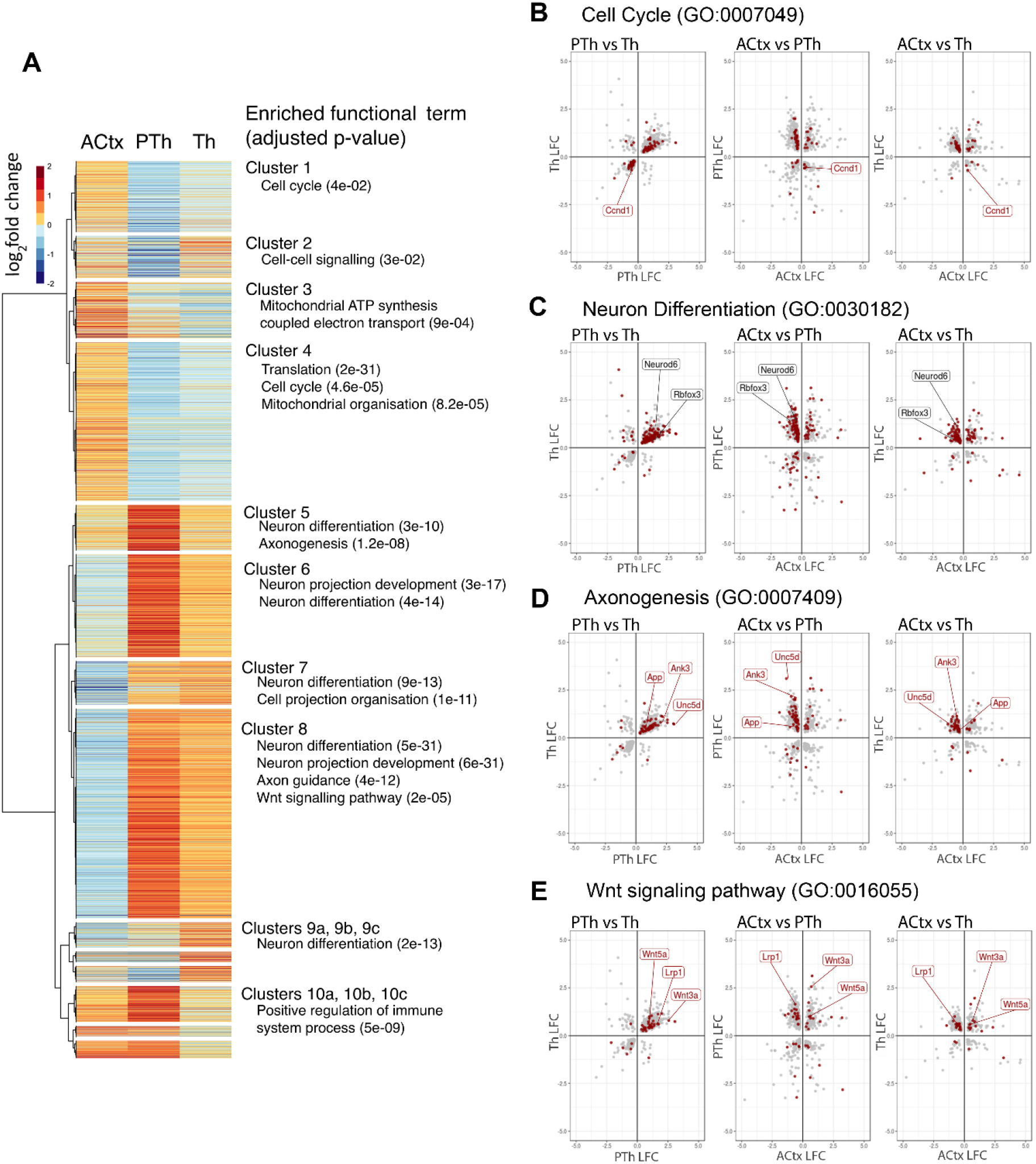
Pax6 deletion has opposite effects on the transcriptomes of cortical and diencephalic (thalamic and prethalamic) cells. (A) Hierarchical clustering of all genes that showed significant differential expression in at least one tissue according to the direction and magnitude of their log_2_ fold change (LFC) across all three tissues. The dendrogram is cut to generate 14 clusters and enriched GO functional terms are listed against these clusters. (B-E) LFCs of all genes showing significant differential expression (adjusted p<0.05) in PTh and Th against each other, in ACtx and PTh against each other and in ACtx and Th against each other. Genes associated with particular GO terms (“cell cycle”, “neuron differentiation”, “axonogenesis” and “Wnt signalling pathway”) are marked on the plots (red). Linked to Supplementary File 6.

Finally, we plotted the LFCs of all genes showing significant differential expression (adjusted p<0.05) in (i) PTh and Th against each other, (ii) ACtx and PTh against each other and (iii) ACtx and Th against each other (Figure 3B-E). Genes associated with particular GO terms (“cell cycle”, “neuron differentiation”, “axonogenesis” and “Wnt signalling pathway”) are marked on the graphs (in red). The LFCs of most genes associated with these and similar GO terms were directly related in Th and PTh (first column of graphs in Figure 3B-E). However, the LFCs of large proportions of these genes were inversely related when Th or PTh were compared to ACtx (second and third columns of graphs in Figure 3B-E). Selected genes associated with each function are highlighted in the graphs. For example, the cell cycle gene *Ccnd1* (*CyclinD1*) was significantly downregulated in both Th and PTh but significantly upregulated in ACtx (Figure 3B). On the other hand, genes associated with differentiation such as *Neurod6* and *Rbfox3* (also commonly known as *NeuN*) were significantly upregulated in Th and PTh but significantly downregulated in ACtx (Figure 3C). To identify in these graphs any individual gene or group of genes associated with any GO term, visit *LFC-GO plots* at https://pricegroup.sbms.mvm.ed.ac.uk/Pax6_diencephalon/.

These analyses showed that Pax6 deletion caused many genes to alter their expression levels in opposite directions in cortical versus diencephalic regions. They indicate that many of these oppositely regulated genes are associated with cell proliferation and differentiation. Moreover, they suggest that Pax6 loss is likely to promote proliferation over differentiation in cortex but to have the opposite effect in both diencephalic regions.

### Pax6 has opposite effects on neurogenesis in cortex and diencephalon

We then tested this prediction directly in histological sections of embryonic brain. As before, we created *CAG^CreER^ Pax6* cKOs and littermate controls by giving tamoxifen at E9.5. We carried out the analysis at three different ages for all tissues (E11.5, E12.5 and E13.5), thereby allowing for the fact that diencephalic tissues develop slightly ahead of telencephalic tissues, with neurogenesis starting by E11.5 in the former (Suzuki-Hirano et al., 2011) and by E12.5 in the latter. We injected a single pulse of the S-phase marker bromodeoxyuridine (BrdU) 24h before collecting tissue for analysis using markers of proliferating cells (Ki67) and differentiating neurons (Tuj1). We counted from two regions of ACtx, one lateral and one medial, and one region each of Th and Pth (Figure 4A-D’). We measured the proportions of BrdU+ cells that (i) remained proliferative (BrdU+, Ki67+, Tuj1-; white arrows in Figure 4E,F); (ii) had started to differentiate (BrdU+, Ki67-, Tuj1+; red arrows in Figure 4E,F); (iii) were in an intermediate state (BrdU+, Ki67-, Tuj1-; green arrows in Figure 4E) in ACtx, Th and PTh of control and *Pax6* cKO embryos.

**Figure 4.**
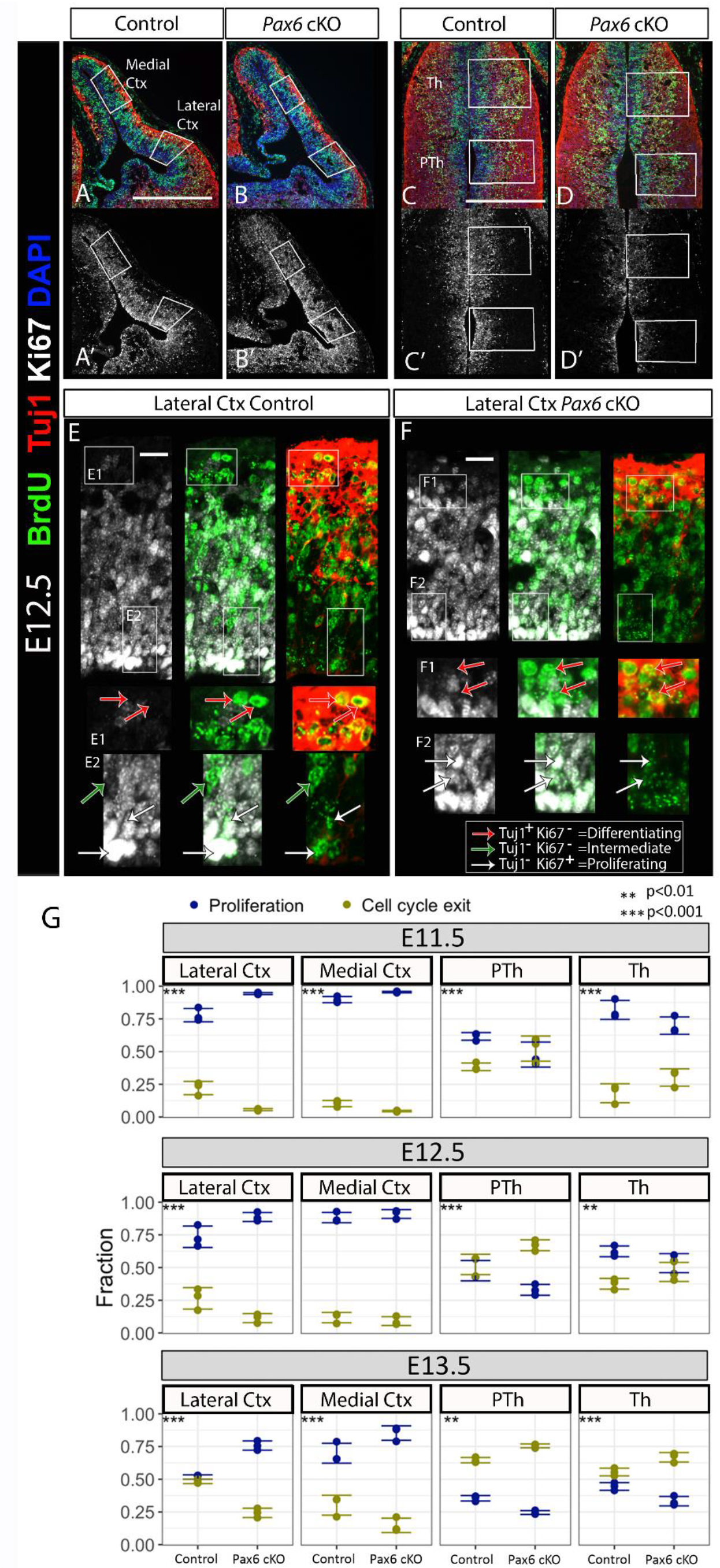
The effect of Pax6 deletion on proliferation and differentiation in cortical versus diencephalic neurons is opposite and not age-dependent. (A-D) Immunohistochemistry for BrdU and Tuj1 on E12.5 cortical (A,B) and diencephalic (C,D) tissues. Boxes show the areas of quantification. Scale bars: 0.25mm (A’-D’) Same sections than A-D, channel showing Ki67 staining has been separated here for a clearer visualization. (E,F) High power details of E12.5 control (E) and *CAG^CreER^ Pax6* cKO (F) lateral cortex showing separate channels. Zoomed boxed areas (E1,E2,F1,F2) show examples of cells attributed to each category in our quantifications (red arrows: differentiated cells, Brdu^+^/Tuj1^+^/Ki67^-^; green arrows: intermediate G0 state, Brdu^+^/Tuj1^-^/Ki67^-^; white arrows: proliferative cells, Brdu^+^/Tuj^-^/Ki67^+^). Scale bars: 10μm (G) Analysis of the quantification of proliferation and cell cycle exit indexes in all ages (E11.5, E12.5, E13.5), tissues (lateral cortex, medial cortex, thalamus and prethalamus) and genotypes (controls and *CAG^CreER^ Pax6* cKO) showed that Pax6 loss produces an increase in proliferation and decrease in cell cycle exit in cortical tissues but the opposite is observed in diencephalic (thalamus and prethalamus) tissues. The data from all ages, regions and genotypes were fitted to a generalized linear mixed model to test the effects of Pax6 inactivation depending on age and tissue (n=3). ANOVA was used with Tukey’s method for multiple pairwise comparisons to obtain test statistics for contrasts in question. LAT CTX, lateral cortex; MED CTX, medial cortex; PTHAL, prethalamus: THAL, thalamus. Linked to Figure 4-figure supplement 1.

Very few cells were BrdU+, Ki67- and Tuj1- in any condition and their numbers changed in proportion to the fraction of differentiated cells (Figure 4-figure supplement 1). Most likely these cells had only recently entered G0 en route to undergoing differentiation. Since there was no evidence that Pax6 deletion caused their numbers to increase, we decided to combine all BrdU+, Ki67-cells into one group representing those that had exited the cell cycle to differentiate.

The data from all ages, regions and genotypes were fitted to a generalized linear mixed model to test the effects of Pax6 inactivation depending on age and tissue (Figure 4G). In the cortex, the proportions of mitotic (BrdU+) cells that remained proliferative (Ki67+) after 24h were significantly higher in cKOs than in controls and the proportions that had exited the cell cycle were significantly lower in cKOs, with the sole exception of the medial cortex at E12.5 where we detected no significant effect. In both Th and PTh, effects of Pax6 loss were opposite to those in the cortex. In both regions at all ages, the proportions of mitotic cells that remained proliferative after 24h were significantly lower in cKOs than in controls and the proportions that had exited the cell cycle were significantly higher.

These data also allowed us to compare the states of maturation of tissues of different ages in control animals. In E13.5 lateral cortex, for example, ^~^45% of mitotic cells had exited the cell cycle during the last 24h. Similar rates of exit are found at E12.5 in Th and are likely to be reached between E11.5 and E12.5 in PTh. This validates our premise that the range of ages included in these experiments allows comparison of the effects of Pax6 at equivalent developmental stages in the three tissues. The effect observed is therefore tissue-specific and not age-dependent. The results agree with our prediction from the RNA-seq experiments that the effects of Pax6 on the balance between proliferation and differentiation are opposite in cortex versus diencephalon.

### Interregional variation in Pax6 levels and exon usage do not correlate closely with its function

We next considered possible mechanisms by which Pax6 might regulate the balance between proliferation and differentiation in opposite directions in cortical and diencephalic regions. The first possibility was that regional variation in attributes of the transcription factor itself cause it to operate differently in cortex and diencephalon. Variation in Pax6’s level and the ratio between its two major splice variants might modify its function (Haubst et al., 2004; Manuel et al., 2006).

Regional variation in levels of Pax6 protein in normal forebrain did not correlate with the effects of Pax6 on the balance of proliferation versus differentiation. Levels of Pax6 are highest in PTh and ACtx, which show opposite changes in proliferation versus differentiation following Pax6 deletion, and relatively low in Th, which shows changes in the same direction as PTh (Figure 1A-D). Regarding alternative splicing, previous studies have suggested that the two major isoforms, Pax6 and Pax6(5a) - the latter contains an additional 14 amino acids inserted into the paired domain - affect gene expression differently (Chauhan et al., 2004; Haubst et al., 2004; Pinson et al., 2005). We used our RNA-seq data to compare exon 5/5a usage in control ACtx, Th and PTh (Figure 5). In each sample, we measured average coverage across exons 5 and 5a and found that the only significant difference was a ^~^2 fold difference in the 5/5a ratio between ACtx and PTh, with the ACtx having higher relative counts for exon 5a and therefore a lower 5/5a ratio. As with Pax6 levels, variation in relative levels of exon 5/5a usage did not correlate closely with the effects of Pax6 on the balance of proliferation versus differentiation. Pax6 is pro-proliferative in both Th and PTh and the opposite in ACtx, whereas the exon 5/5a ratio differed significantly between ACtx and only one of the diencephalic regions. Whereas regional differences in the exon 5/5a ratio might play an important role, on their own they are unlikely to explain regional differences in Pax6 action on the balance of proliferation and differentiation. We went on to consider further possible mechanisms.

**Figure 5.**
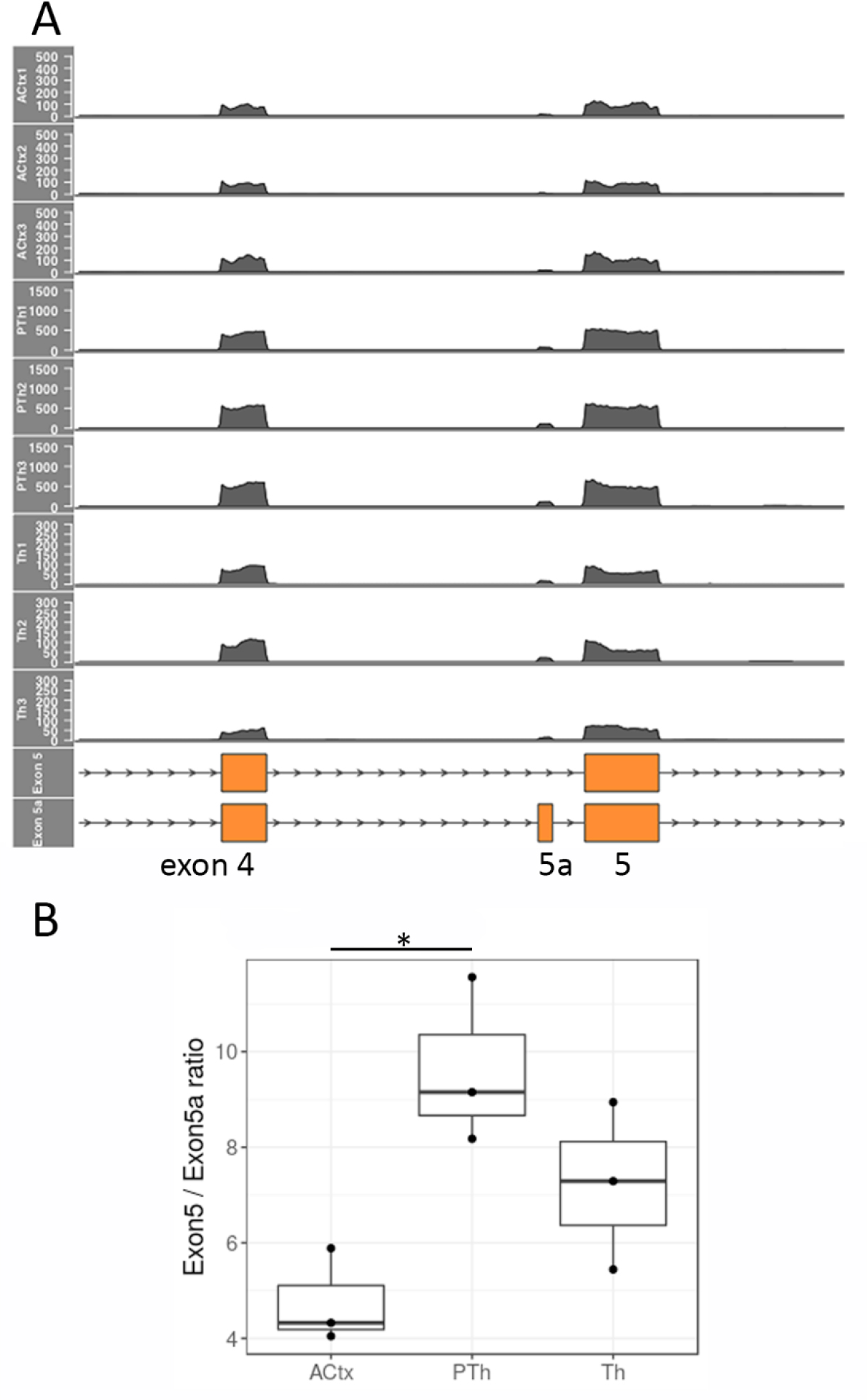
*Pax6*/*Pax6(5a)* ratio in ACtx, Th and PTh in control embryos. (A) Counts per base read coverage of exons 4, 5a and 5 of *Pax6* from each sample. (B) Ratios between the average coverage per base in exons 5 and 5a for each sample superimposed on box and whisker plots. ANOVA followed by post hoc Tukey comparison showed a significant difference between the ratios in ACtx and PTh (p=0.02).

### Pax6 loss causes canonical Wnt signaling deregulation

We then considered the possibility that Pax6 might affect the balance between proliferation and differentiation in opposite directions in cortex and diencephalon because of its differential effects on intercellular signaling. We examined Wnt signaling, since results above identified it as a GO function that is regulated differentially between cortical and diencephalic regions (Figure 3).

A number of genes involved in Wnt signaling were commonly deregulated across all three tissues, sometimes in different directions but sometimes in the same direction. *Lrp1*, which encodes Low Density Lipoprotein Receptor-1, a potential negative regulator of canonical Wnt signalling (Willnow et al., 2007; Zilberberg et al., 2004), was upregulated in Th and PTh but downregulated in ACtx (Figure 3E). While these changes correlated with regional differences in the effects of Pax6 on the balance of proliferation versus differentiation, others did not. Wnt3a and Wnt5a were upregulated across all three regions. The Wnt antagonists *Sfrp2* and *Dkk3* were strongly downregulated in prethalamic progenitors but were unaffected in Th, where their levels are normally very low (Figure 6A-G). *Sfrp2* and *Dkk3* were significantly downregulated in ACtx, although their expression domains are limited mainly to extremely lateral cells around the pallial-subpallial boundary and medially at the cortical hem respectively (Figure 6B,C,F,G; Kim et al., 2001).

**Figure 6.**
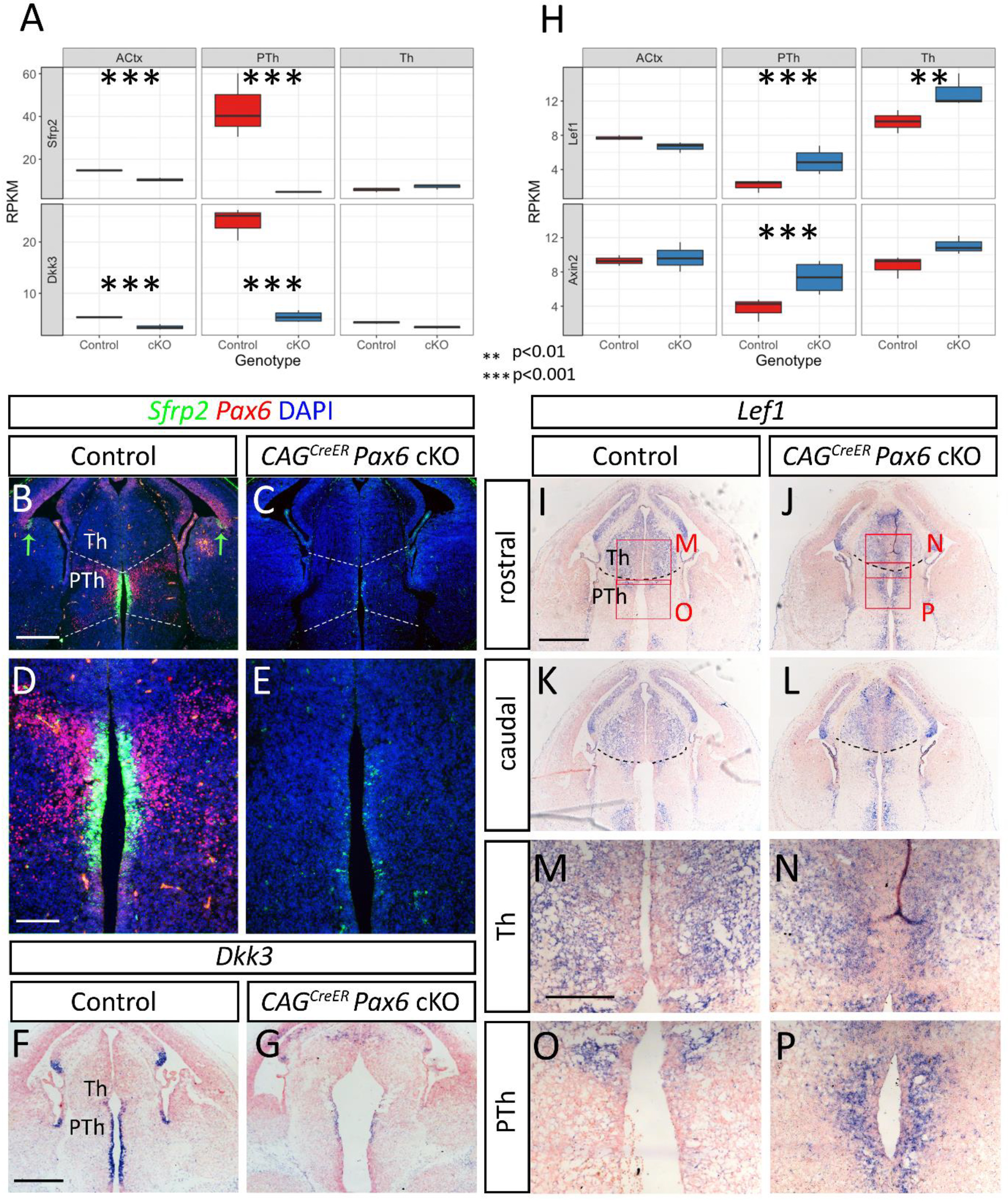
Pax6-loss-induced changes in canonical Wnt signalling in the forebrain. (A,H) Box and whisker plots showing data from RNA-seq of reads per kilobase of transcript per million mapped reads (RPKM) for *Sfrp2*, *Dkk3*, *Lef1* and *Axin2* in ACtx, PTh and Th from control and *CAG^CreER^ Pax6* cKOs. Significance was tested by ANOVA followed by post hoc Tukey comparison. (B-G,I-P) In situ hybridizations for *Sfrp2*, *Dkk3* and *Lef1* and immunohistochemistry for Pax6 in E13.5 control and *CAG^CreER^ Pax6* cKOs. Green arrows in B indicate *Sfrp2*+ cells at the pallial-subpallial boundary. Scale bars: B,C,F,G,I-L, 0.25mm; D,E,M-P, 0.1mm.

To see if the changes in expression of individual Wnt pathway genes produced a net effect that correlated with regional differences in the effects of Pax6 on the balance of proliferation versus differentiation, we examined the expression of two bona-fide readouts of the Wnt canonical pathway, *Lef1* and *Axin2* (Figure 6H-P). Both genes were significantly upregulated in PTh and *Lef1* was upregulated in Th, but neither gene was significantly altered in ACtx.

These analyses suggest that the effects of Pax6 loss on canonical Wnt signaling are largely restricted to the diencephalon, where the effects were greatest in PTh. While these changes would likely affect the behaviors of diencephalic progenitors, they do not provide a clear explanation for the effect of Pax6 loss on cortical progenitors. We went on, therefore, to explore other factors that might be important for explaining the nature of Pax6’s cortical effect.

### Foxg1-null telencephalon develops a diencephalon-like profile of cell cycle gene expression

We next examined the possibility that Pax6 operates differently in cortex because another transcription factor in this tissue moves the transcriptomes of cortical cells away from those of diencephalic cells, thereby altering the context within which Pax6 operates and so modifying its actions. We postulated that such a factor should be present in cortex and not diencephalon and that it should not be regulated by Pax6. Transcripts for transcription factor Foxg1 showed the highest log_2_ fold enrichment between ACtx and Th (11.67) and the third highest between ACtx and PTh (9.57) (Supplementary File 1; Figure 1-figure supplement 2Y). Foxg1 is co-expressed with Pax6 by cortical progenitors and is an important positive regulator of their cell cycles (Martynoga et al., 2005; Vezzali et al., 2016; Xuan et al., 1995; Yip et al., 2012). Our RNA-seq data showed that its cortical expression is only marginally downregulated following Pax6 removal (LFC=-0.207; adjusted p=0.0190), a change that was not visible in cortical sections (Figure 7A,B). Levels of Foxg1 protein and its nuclear distribution in cortical cells appeared unaltered by Pax6 loss (Figure 7D-F). We hypothesized, therefore, that Foxg1 might modify the molecular context within which Pax6 operates and therefore its actions.

**Figure 7.**
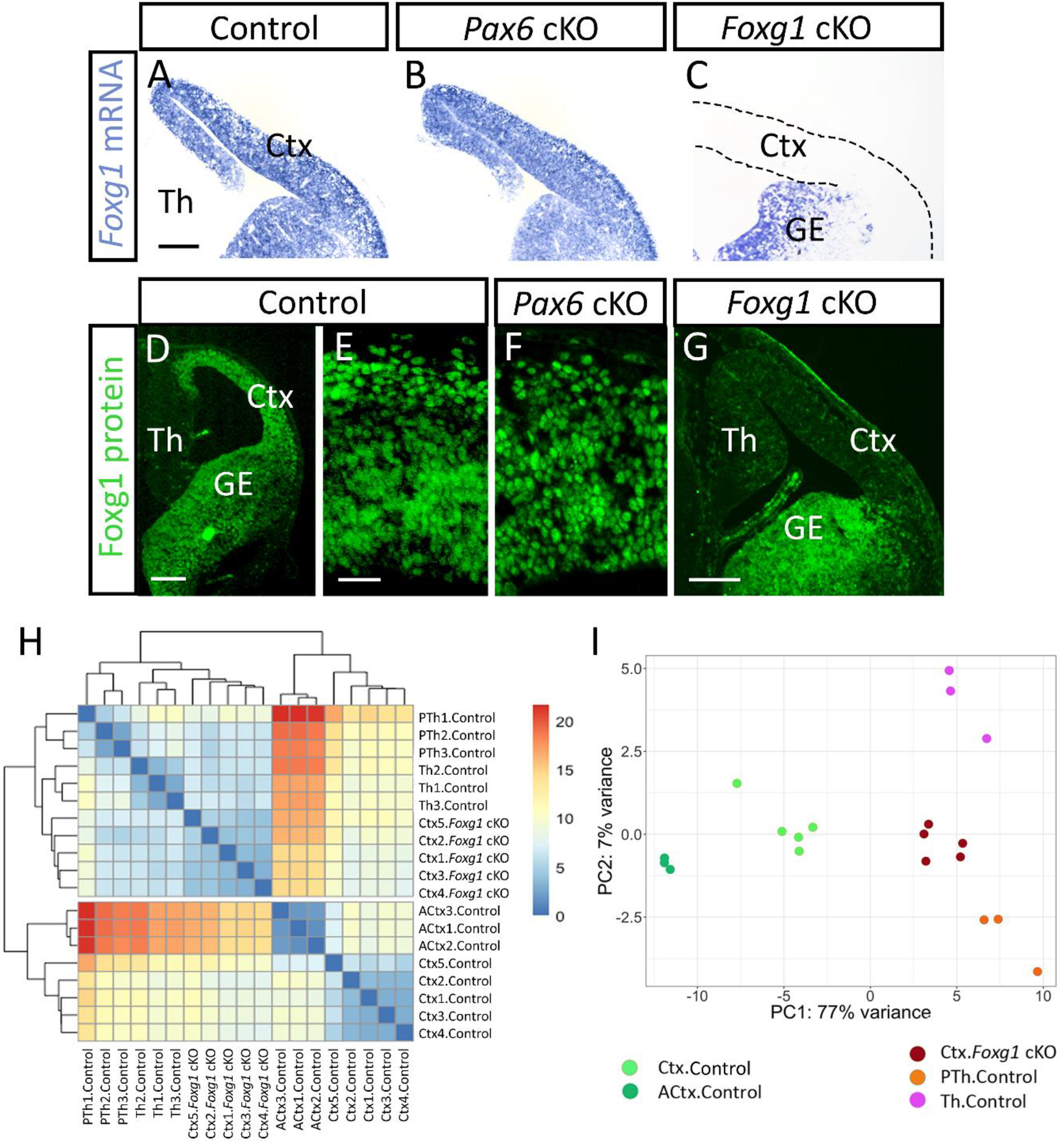
Foxg1 deletion in the embryonic cortex makes its profile of expression of cell cycle genes diencephalon-like. (A-C) In situ hybridizations showing *Foxg1* telencephalic expression in control, *Emx1^CreER^ Pax6* cKOs and *Emx1^CreER^ Foxg1* cKOs at E13.5. GE, ganglionic eminence. Scale bar: 0.2mm. (D-G) Immunohistochemistry showing Foxg1 protein expression in (D) control telencephalon, (E) control cortex, (F) *Emx1^CreER^ Pax6* cKO cortex and (G) *Emx1^CreER^ Foxg1* cKO telencephalon at E13.5. Scale bars: D,G, 0.2mm; E,F, 0.05mm. (H) Heatmap of hierarchical clustering of RNA-seq data on genes annotated by the “Cell Cycle” GO term (GO:0007049) that were significantly deregulated in the *Foxg1* cKO. Data are from samples of control cortex, control ACtx, control PTh, control Th and *Emx1^CreER^ Foxg1* cKO cortex. (H) Principal component (PC) analysis on the same RNA-seq data as in (H). Linked to Figure 7-figure supplement 1, Figure 7-figure supplement 2 and Supplementary files 7,8 and 9.

We first tested whether differences between the transcriptomes of cortical and diencephalic cells are created by Foxg1’s presence in the cortex. We used RNA-seq to test whether Foxg1 deletion from the cortex moved its profile of gene expression towards that of diencephalic tissues. We gave tamoxifen at E9.5 to induce cortex-specific deletion of *Foxg1* (*Emx1^CreER^*; *Foxg1^fl/fl^*, referred to here as *Foxg1* cKO; the *Foxg1^fl^* allele was from Miyoshi and Fishell, 2012 which caused loss of Foxg1 from almost all cortical cells by E13.5 (Figure 7C,G). We carried out RNA-seq in E13.5 control (*Emx1^CreER^*; *Foxg1^fl/^*^+^) and *Foxg1* cKO cortices (in these experiments the entire cortex was used). Five replicate samples for controls and cKOs from independent litters were sequenced. The raw data can be obtained from the European Nucleotide Archive (www.ebi.ac.uk/ena; Project 10900, ENA accession numbers PRJEB21349, ERP023591). PCA showed that data from individual samples clustered by genotype along the axis of greatest variation (Figure 7-figure supplement 1A).

Supplementary File 7 lists all genes with significant differential expression in *Foxg1* cKOs (adjusted p<0.05) and the numbers of upregulated and downregulated genes are shown in Figure 7-figure supplement 1B,C. The list of downregulated genes showed high enrichment for functional terms related to cell cycle and mitosis (Supplementary File 8), consistent with the fact that Foxg1 is a regulator of cortical proliferation (Martynoga et al., 2005; Xuan et al., 1995). This effect is unlikely to be mediated by an effect of Foxg1 on Pax6 levels or alternative splicing. Pax6 levels were only marginally decreased at the RNA level (LFC = −0.18; adjusted p=0.007), Pax6 protein remained expressed in a normal pattern (Figure 7-figure supplement 1D) and there was no significant effect of Foxg1 deletion on the relative usage of Pax6’s exons 5 and 5a (Figure 7-figure supplement 1E,F). We conclude that both Foxg1 and Pax6 operate independently with no significant effects of either gene on expression of the other.

To find out whether Foxg1 modifies the context within which Pax6 operates by shifting it towards that in diencephalic tissues, we compared RNA-seq data from the cortex of *Foxg1* cKOs with those from control cortex, Th and PTh. Two sets of control cortical data were used: from the five samples obtained as controls for the *Foxg1* cKOs and the three samples from ACtx obtained as controls for the *CAG^CreER^ Pax6* cKOs. Control thalamic and prethalamic data were from the samples obtained as controls for the *CAG^CreER^ Pax6* cKOs. We carried out distance-based hierarchical clustering including specifically those genes annotated by the “Cell Cycle” GO term that were significantly deregulated in the *Foxg1* cKO. All 8 control cortical samples clustered together and separated not only from control diencephalic samples but also from *Foxg1*-null cortical samples (Figure 7H; Supplementary File 9 lists the genes included in this analysis). Control diencephalic samples were clustered with *Foxg1*-null cortical samples. Only at the third level in the dendrogram did *Foxg1* cKO cortical samples separate from Th control samples. PCA using the same set of genes revealed the same trend: *Foxg1*-null cortical samples, but not control cortical samples, clustered with samples from control Th and PTh along the axis of maximum variation, which represented 77% of the variance (Figure 7I). We repeated the same clustering experiments with sets of genes annotated by other GO terms but none showed the same separation as observed for cell cycle genes (examples are shown in Figure 7-figure supplement 2).

These results suggest that the deletion of Foxg1 from the cortex shifts the profile of expression of genes whose actions are related to the cell cycle closer to that of normal diencephalic tissues. We then went on to test whether this shift alters the way in which cortical cells *respond* to Pax6 by asking whether Pax6 deletion from *Foxg1^-/-^* mutant telencephalon causes a response closer to that observed when Pax6 is lost from the wild type diencephalon.

### Foxg1 alters Pax6’s effect on expression of an important cell cycle regulator

Before testing whether the presence of Foxg1 in the cortex modifies the effects of Pax6 on the balance between proliferation and differentiation, we sought molecular evidence for whether this was likely. We first focussed on those genes that were regulated by both Pax6 and Foxg1 obtained from the intersection of our RNA-seq data from *Pax6* cKOs and *Foxg1* cKOs. We identified 678 genes whose cortical expression was regulated by both Pax6 and Foxg1. These genes are listed in Supplementary File 10 and their average LFCs in *Pax6* cKO and *Foxg1* cKO are plotted against each other in Figure 8A. Pax6 and Foxg1 regulate many of these genes in the same direction but many in opposite directions. We focussed on the latter group. We argued that this group would be most likely to include genes whose expression was affected because Foxg1 reversed Pax6’s actions on them.

**Figure 8.**
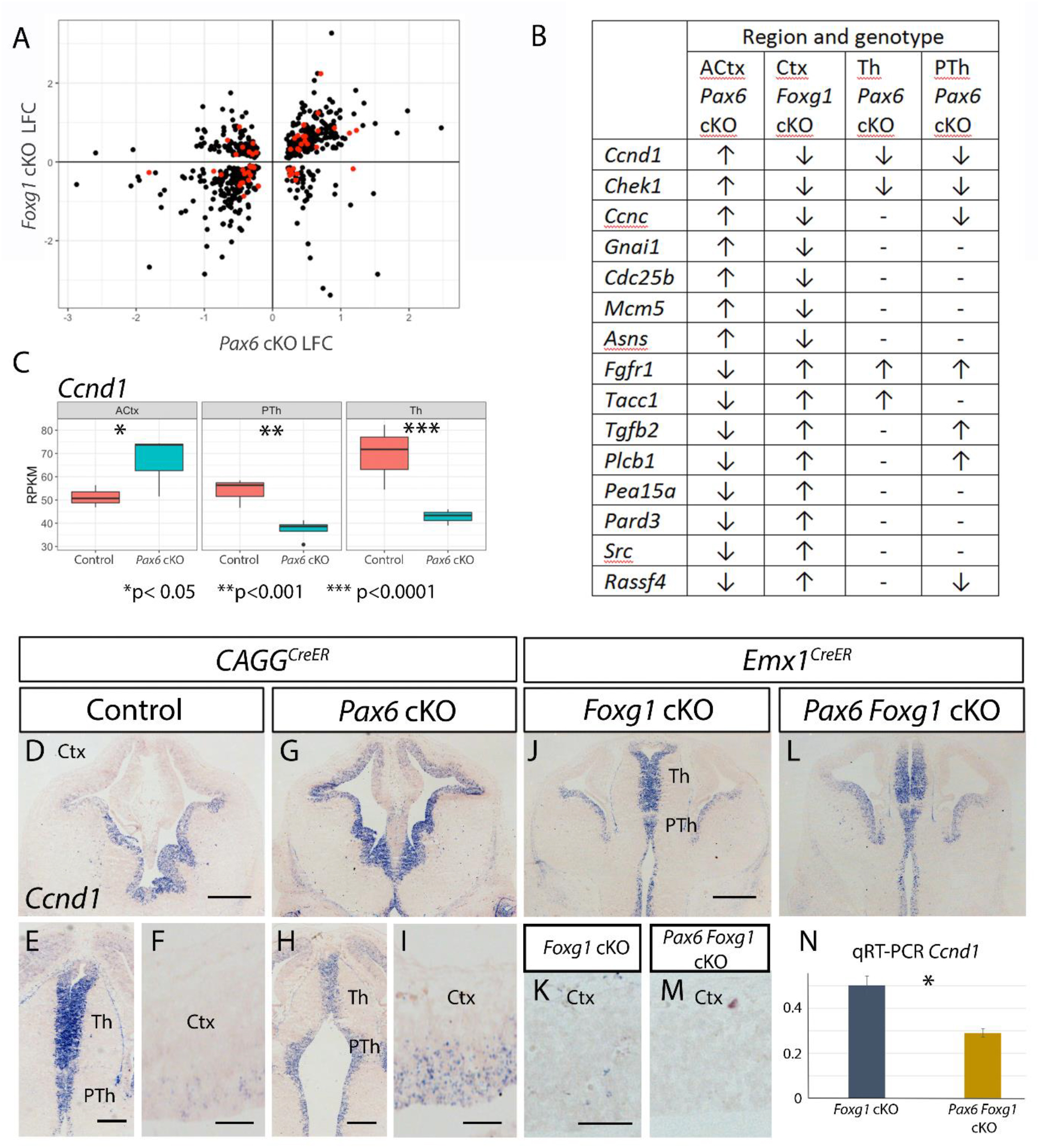
Foxg1 deletion in the embryonic cortex reverses effects of Pax6 deletion. (A) Log_2_ fold changes (LFCs) of all genes showing significant regulation by both Pax6 (data from *CAG^CreER^ Pax6* cKOs) and Foxg1 (data from *Emx1^CreER^ Foxg1* cKOs). Red dots mark cell cycle associated genes. (B) Genes associated with the cell cycle regulated in opposite directions in *CAG^CreER^ Pax6* cKOs versus *Foxg1* cKOs; any significant changes following Pax6 removal from Th or PTh are indicated. (C) Box and whisker plots of RPKM for *Ccnd1* in ACtx, PTh and Th from E13.5 control and *CAG^CreER^ Pax6* cKOs. Significance was tested by ANOVA followed by post hoc Tukey comparison. (D-I) In situ hybridizations showing expression of *Ccnd1* in the forebrains of E13.5 controls and *CAG^CreER^ Pax6* cKOs. Scale bars: D,G, 0.25mm; E,H, 0.2mm; F,I, 0.05mm. (J-M) In situ hybridizations showing expression of *Ccnd1* in the forebrains of E13.5 *Emx1^CreER^ Foxg1* cKOs and *Emx1^CreER^ Foxg1 Pax6* cKOs. Scale bars: J,L, 0.25mm; K,M, 0.05mm. (N) Quantitative RT-PCR for *Ccnd1* in E13.5 cortex of *Emx1^CreER^ Foxg1* cKOs and *Emx1^CreER^ Foxg1 Pax6* double cKOs. Means ± sems; n=3 for each genotype; p<0.05 Student’s t-test. Linked to Supplementary file 10.

Of the genes whose expression was deregulated in opposite directions by cortical deletion of Pax6 or Foxg1, 15 had GO annotations relating them with cell cycle control (red dots in Figure 8A). These genes are listed in Figure 8B along with the direction of their expression change when either Pax6 or Foxg1 was removed from each region. Seven of these genes showed directions of change in *CAG^CerER^ Pax6* cKOs that were opposite in ACtx versus Th and/ or PTh. These gene expression changes, individually or collectively, might make a significant contribution to changing the balance between proliferation and differentiation in opposite directions in cortex versus the diencephalon after Pax6 deletion.

One of these genes was the cell cycle regulator *Ccnd1*, which we took as an exemplar of the principles outlined above. In our RNA-seq data, *Ccnd1* showed significant Pax6-loss-induced differential expression in opposite directions in cortex (upregulated) and diencephalon (downregulated in both Th and PTh) (Figure 8C). These changes can be seen with in situ hybridization in Figure 8D-I. Ccnd1 is a G1 phase cyclin which promotes entry into S-phase and hence cell cycle progression over cell cycle exit in many systems and cell types.

Importantly, several previous studies have shown that altering its expression levels in the embryonic cortex alters the balance between proliferation and differentiation (Artegiani et al., 2011; Ferguson et al., 2000; Kollmann and Sexl, 2013; Lange et al., 2009; Matsushime et al., 1994; Morgan, 1997; Pilaz et al., 2009; Zerjatke et al., 2017). In the light of these findings, we studied changes in *Ccnd1* expression further since they were likely to be good predictors of cellular responses to Pax6 removal in different contexts.

To test whether Foxg1 affects the actions of Pax6 on *Ccnd1* expression, we tested the effect of deleting Pax6 from *Foxg1*-null cortex. We generated *Emx1^CreER^*;*Foxg1^fl/fl^* and *Emx1^CreER^*;*Foxg1^fl/fl^*;*Pax6^fl/fl^* embryos and induced deletion of *Foxg1* or both *Pax6* and *Foxg1* by tamoxifen administration at E9.5 to produce embryos referred to here as *Foxg1* cKOs or *Pax6*;*Foxg1* dcKOs (double cKOs) respectively. Both Pax6 and Foxg1 protein were absent from all but the most lateral cells of the cortex, where *Emx1* is not expressed, by E13.5 (Figures 6C,G, 8I,I’; see also Mi et al., 2013). Whereas *Ccdn1* expression was significantly reduced in *Foxg1* cKO cortex (Supplementary File 7), levels were even lower following Pax6 co-deletion (Figure 8J-N). This Pax6-loss-induced change was opposite to that which occurs following Pax6 removal from cortex that expresses Foxg1. It was in the same direction as occurs in Th and PTh following Pax6 loss.

Altogether, our results increased the likelihood that Pax6’s effects on the balance of proliferation versus differentiation in the cortex are Foxg1-dependent. We went on to test the hypothesis that in cortex lacking Foxg1, Pax6’s effects would be more like those normally seen in the diencephalon.

### Absence of Foxg1 alters the cellular response of cortex to Pax6 removal

We first counted the proportions of cortical cells that were proliferative (Ki67+) in E13.5 control, *Pax6* cKO, *Foxg1* cKO and *Pax6*;*Foxg1* dcKO embryos (Figure 9A-E). We found that, in contrast to the consequences of removing Pax6 from normal Foxg1-expressing cortex (Figure 4; Figure 9A,B), removing Pax6 from *Foxg1^-/-^* mutant cortex caused a significant reduction in the proportion of proliferating cells (Figure 9E; Students t-test, p= 0.02). This response was opposite to that of Foxg1-expressing cortex and in the same direction as that of Th and PTh (Figure 4).

**Figure 9.**
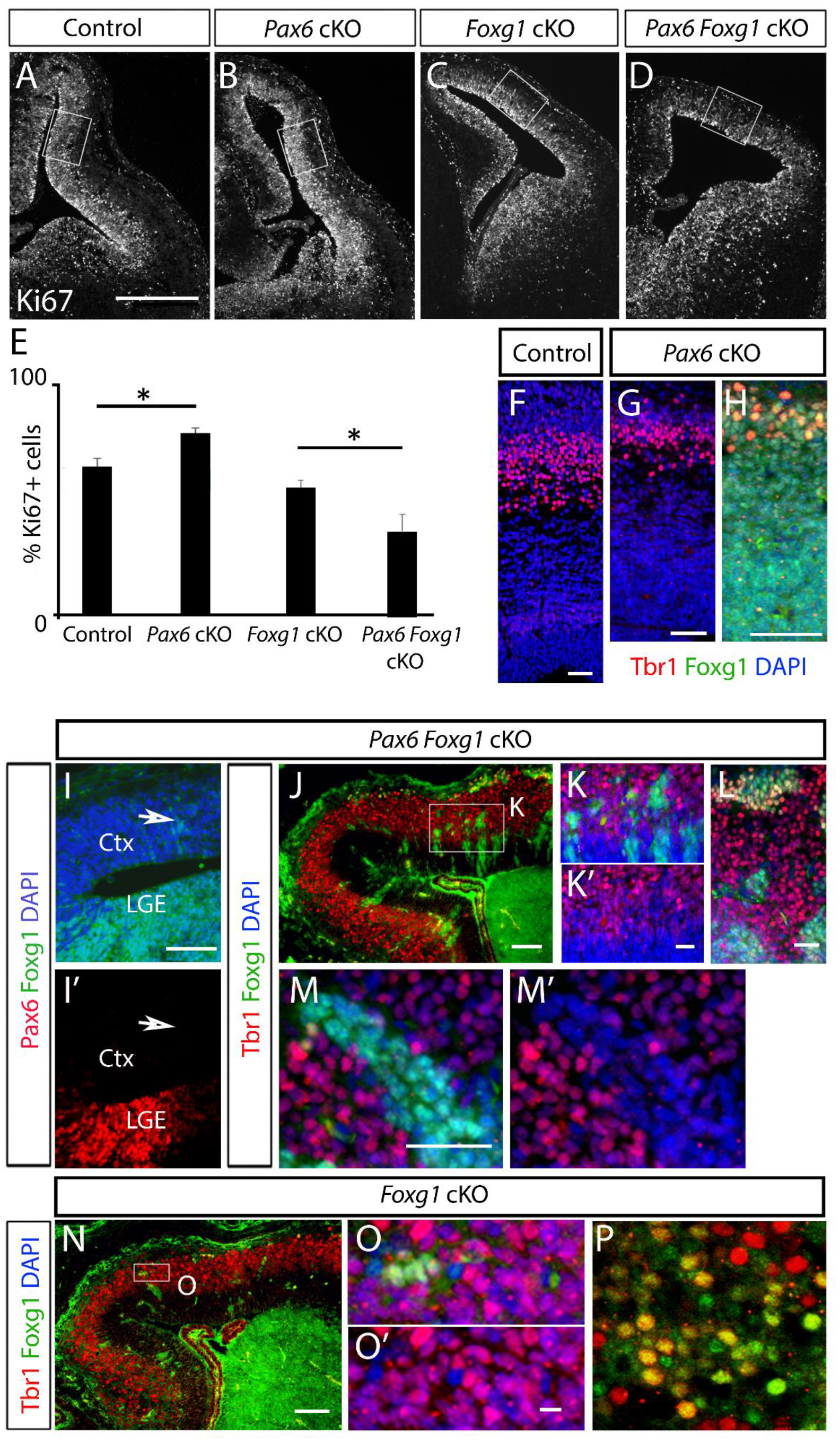
Foxg1 deletion in the embryonic cortex reverses the effects of Pax6 deletion on the balance of proliferation vs differentiation. (A-D) Immunohistochemistry of Ki67+ cells in the cortex of E13.5 controls, *CAG^CreER^ Pax6* cKOs, *Emx1^CreER^ Foxg1* cKOs and *Emx1^CreER^ Foxg1 Pax6* cKOs. Boxes show the areas selected for quantification. Scale bar: 0.25mm. (E) Quantifications of Ki67+ cells in the cortex of E13.5 controls, *CAG^CreER^ Pax6* cKOs, *Emx1^CreER^ Foxg1* cKOs and *Emx1^CreER^ Foxg1 Pax6* cKOs. Means ± sems; n=3 for each genotype; p<0.05 Student’s t-test. (F-H) Immunohistochemistry for Tbr1 and Foxg1 in control and *CAG^CreER^ Pax6* cKO at E15.5. Scale bars, 0.05mm. (I,I’) Immunohistochemistry for Pax6 and Foxg1 on the same section of E13.5 *Emx1^CreER^ Foxg1 Pax6* double cKO telencephalon. Arrow shows that small numbers of cells in the cortex retained Foxg1 but not Pax6. Scale bars: 0.2mm. (J-M’) Immunohistochemistry for Foxg1 and Tbr1 on E15.5 *Emx1^CreER^ Foxg1 Pax6* cKOs. Box in J is shown in K,K’. Scale bars: J, 0.05mm; K-M’, 0.025mm. (N-P) Immunohistochemistry for Foxg1 and Tbr1 on E15.5 *Emx1^CreER^ Foxg1* cKOs. Box in N is shown in O,O’. Scale bars: N, 0.05mm; O-P, 0.01mm.

We then tested the effect of Pax6 deletion from *Foxg1*-null cortex on the production of Tbr1-expressing cells. Tbr1 marks cortical cells as they transition from a proliferative progenitor state to a postmitotic neuronal state (Englund et al., 2005). In our RNA-seq data, *Tbr1* levels were significantly reduced in cortical *Pax6* cKOs (Supplementary File 2), fitting with the shift to proliferation. This was also clear with immunohistochemistry (Figure 9F-H). Here we compared the effects of Pax6 removal from *Foxg1*-null versus Foxg1-expressing cortical cells on their production of Tbr1+ cells at E15.5.

We observed that in some E13.5 and E15.5 *Foxg1* cKOs and *Pax6*;*Foxg1* dcKOs, small clusters of cells continued to express Foxg1 protein. These clusters comprised only a few cells in the ventricular zone at E13.5 (Figure 9I) but were larger by E15.5 (Figure 9J-P), as expected since their failure to delete *Foxg1* would enhance their proliferation relative to *Foxg1^-/-^* cells. Foxg1+ clusters in *Pax6*;*Foxg1* dcKOs did not express Pax6 (arrows in Figure 9I,I’), suggesting that the floxed *Pax6* allele deleted more efficiently than the floxed *Foxg1* allele. This mosaicism allowed us to compare how the presence or absence of Foxg1 in neighbouring cortical cells affected their response to Pax6-deletion.

In clusters that retained Foxg1 expression in E15.5 *Pax6*;*Foxg1* dcKOs (i.e. single *Pax6* cKO cells), only a small percentage of cells expressed Tbr1 (12%±3sd). These had reached the outermost layer of the cortical plate (Figure 9J-M), agreeing closely with the pattern and proportions of Tbr1-expressing cells (11%±2sd) seen in E15.5 *Pax6* single cKO embryos (Figure 9G,H). On the other hand, the majority (59%±7sd) of cells that were null for both *Pax6* and *Foxg1* in dcKOs expressed Tbr1, indicating that neuronal differentiation was much more advanced in this population. To confirm that the differential outcome between Foxg1-positive and Foxg1-negative cells was a consequence of Pax6 removal, we compared clusters of Foxg1-positive and Foxg1-negative cells in *Foxg1* single cKOs, in which *Pax6* was normal. In these embryos, large proportions of both Foxg1-positive and Foxg1-negative cells throughout the cortical plate were Tbr1-positive (Figure 9N-P).

These results indicate that whereas Pax6 normally promotes differentiation over proliferation in embryonic cortex, this effect is Foxg1 dependent. If Foxg1 is absent, Pax6 promotes cortical cell cycle progression and Pax6 removal is anti-proliferative and pro-differentiative.

## Discussion

Previous studies reported that Pax6 has very different effects on the transcriptomes of forebrain cells and highly divergent Pax6-expressing non-neural cell types, namely the lens of the eye and β-cells of the pancreas (Mitchell et al., 2017; Sun et al., 2015; Xie et al., 2013). However, the molecular actions of Pax6 within specific subdivisions of the forebrain have not previously been reported. In this study, we explored the differences between the transcriptomes of embryonic cortical and diencephalic (thalamic and prethalamic) cells in control brains and in brains from which Pax6 had been deleted. There were many more differences between control cortical and diencephalic cells than between thalamic and prethalamic cells. There were also many more differences between control cortical and diencephalic cells than were introduced within either tissue by Pax6 loss. We found substantial differences in the sets of genes whose expression changed in response to deletion of Pax6 in cortex and diencephalon. Indeed, our experiments at both molecular and cellular levels demonstrated that Pax6 regulates the balance between proliferation and differentiation in opposite directions in cortex versus diencephalon. The actions of this key transcription factor are clearly very different in each of these forebrain regions.

We then tested several possible mechanisms to explain the cause of these interregional differences in Pax6’s actions within the forebrain. We found that the presence of Foxg1 in the cortex is a major factor. In vertebrates, the presence of Foxg1 protein is a hallmark of telencephalon (Kumamoto and Hanashima, 2017). Its expression is restricted to the telencephalic anlage in the anterior neural plate as this region folds, closes and expands to form cerebral cortex dorsally and basal ganglia ventrally. It is present in the nuclei of telencephalic progenitors and affects their proliferation, maintaining a normal rate of cell cycle progression and preventing their exit from the cell cycle (Martynoga et al., 2005; Xuan et al., 1995). Its ability to bind DNA is required for it to keep progenitors in a proliferative state (Hanashima et al., 2002).). This requirement is cell autonomous, as evidenced by our previous work on the cortex of *Foxg1^-/-^*;*Foxg1^+/+^* chimeras showing that abnormalities of proliferation occur in *Foxg1*-null cells even if they are surrounded by wild-type cells (Manuel et al., 2011). Foxg1 is highly conserved not only in vertebrates but also in invertebrates that lack a telencephalon (Bredenkamp et al., 2007). In invertebrates, Foxg1 or its homologues are expressed in, and have an instructive role in the development of, cells situated anteriorly in the developing nervous system (Grossniklaus et al., 1994; Pani et al., 2012; Toresson et al., 1998). It is likely that the acquisition of Foxg1 by anterior neural tissue has been an important driver of this region’s evolutionary expansion.

Like Foxg1, Pax6 is highly conserved across the animal kingdom and is essential for normal anterior neural development in both invertebrates and vertebrates (Manuel et al., 2015; Yuan et al., 2016). Unlike Foxg1, however, Pax6 promotes exit from the cell cycle in the mammalian cerebral cortex. On the face of it, this seems paradoxical since it would limit the production of progenitors and hence ultimately the numbers of neurons in a structure whose major defining feature is its rapid evolutionary expansion. It is conceivable that Pax6 actually evolved as a promoter of cell cycle re-entry, a feature that it demonstrates in the diencephalon, but that the superimposition of Foxg1 expression in the telencephalon has had the effect of reversing this activity. In a context where Foxg1 is generating a powerful drive to cell cycle re-entry, Pax6, although possessing the potential to promote proliferation, might have become a brake to Foxg1’s drive, preventing the counterproductive retention of large numbers of cells in a progenitor state. How the expression of Foxg1 leads to a reversal of Pax6’s actions on proliferation is likely to be complex, with some or all of a number of possible mechanisms operating.

The effect of the transcription factor Pax6 on the cell cycle is far from clear. Moreover, its role in different tumor cell types is contradictory. It has been shown to have both oncogenic and tumor suppressor effect depending on the tissue affected (Hegge et al., 2018; Muratovska et al., 2003; Zhang et al., 2015; Zhou et al., 2005). Here we show one mechanism by which this versatile transcription factor can control the cell cycle in opposite ways, depending on the presence or absence of another important transcription factor. This might be important for determining the nature of Pax6’s role in different tumour types.

Pax6 and Foxg1 might converge directly on the same downstream cell cycle genes to regulate their expression. For example, ChIP-seq studies on Pax6 and Foxg1 have shown that both transcription factors can bind genomic regions likely to regulate the expression of cell cycle regulators including *Ccnd1* (Bulstrode et al., 2017; Sun et al., 2015). In the absence of Foxg1, Pax6 binding might activate expression of genes such as *Ccnd1* that promote cell cycle re-entry. Pax6 binding might have an opposite effect if Foxg1 is present since Pax6 binding, although pro-proliferative, might interfere with a potentially more powerful pro-proliferative effect of Foxg1 binding, resulting in an inhibition of cell cycle re-entry. In other words, removal of Pax6 might allow Foxg1 freer rein to drive proliferation. This could occur through direct competition between Pax6 and Foxg1 for binding, a phenomenon observed in other systems (Hong and Wu, 2010; Ilsley et al., 2017; Ngondo-Mbongo et al., 2013; Norton et al., 2017; Wan et al., 2011; Zabet and Adryan, 2013). Another possibility is that, in common with other Fox transcription factors, Foxg1 might be a chromatin remodelling (or pioneer) transcription factor that opens the chromatin to allow proteins such as Pax6 to access sites that they would not otherwise be able to bind, thereby changing its function (Golson and Kaestner, 2016; Iwafuchi-Doi and Zaret, 2016; Magnani et al., 2011). Another possibility is that Foxg1 modifies Pax6’s function by binding to it rather than to DNA.

Pax6 and Foxg1 might not converge directly on the same downstream cell cycle genes but rather they might act on progenitors independently through separate routes. Our data indicate that there are many more differences than similarities in the genes regulated by Pax6 and Foxg1, with both regulating many other transcription factors that have actions on progenitor proliferation. It is possible that these Pax6- and Foxg1-regulated transcription factors then control the activity of more direct cell cycle regulators. Indeed, given the breadth of Pax6’s and Foxg1’s actions, it seems likely that multiple mechanisms will need to be invoked to explain how Pax6 affects the balance of proliferation and differentiation in opposite directions in cortex and diencephalon.

One possibility we considered was that regional variation in an attribute of Pax6 itself, such as its expression level or its exon usage, might cause it to operate differently in cortex and diencephalon. Variation in Pax6’s level and the ratio between its two major splice variants, Pax6 and Pax6(5a), can modify its function (Chauhan et al., 2004; Haubst et al., 2004; Manuel et al., 2006). We found, however, that regional variation in levels of Pax6 protein and in the ratio between exon 5/5a usage did not correlate in a straightforward way with the effects of Pax6 on the balance of proliferation versus differentiation. Previous work has shown that both Pax6 and its rarer variant Pax6(5a) can suppress proliferation and that the level of suppression is dose-dependent across the cortex (Haubst et al., 2004; Manuel et al., 2006; Mi et al., 2013). Levels of Pax6 are, however, very high in PTh, where our new evidence indicates that Pax6 promotes proliferation, an effect that is shared with neighbouring Th, where Pax6 levels are relatively low. It is unclear how the ~2 fold difference in exon 5/5a usage between ACtx and PTh, but not between ACtx and Th, might contribute to the opposite effects of Pax6 on proliferation in ACtx versus both PTh and Th. It is conceivable that the actions of the two forms might oppose each other in the context of prethalamic cells and that higher levels of Pax6 relative Pax6(5a) in PTh might promote proliferation in this region, but at present there is no evidence for or against this possibility.

We also considered the possibility that a differential effect on canonical Wnt signalling might contribute. In eye development, Pax6 directly and positively regulates expression of Wnt inhibitors such as *Sfrp1*, *Sfrp2* and *Dkk1*, thereby suppressing canonical Wnt signalling, with absence of Pax6 leading to aberrant canonical Wnt activity (Machon et al., 2010). In the forebrain, our results indicated that Pax6 loss caused significant upregulation of canonical Wnt signaling (indicated by increased *Lef1* and *Axin2* expression) in diencephalic structures but not in cortex. It appears that Pax6’s effects on canonical Wnt signaling are largely restricted to the diencephalon, where many are striking, such as the almost complete loss of the Wnt antagonist *Sfrp2* from PTh progenitors. While these changes would likely affect the behaviors of diencephalic progenitors, previous studies would predict increased canonical Wnt signaling to promote proliferation (Stolz and Bastians, 2015) whereas we observed the opposite following Pax6 loss from Th and PTh. It seems very likely that multiple synergistic and antagonistic factors combine to generate the tissue-specificity of Pax6’s net cellular effects, but we need a much better understanding of how contributing factors act and interact in the different contexts presented by different brain regions.

In conclusion, we have discovered that Pax6 has radically different molecular and cellular effects on the balance of proliferation and differentiation in two major forebrain regions, the cortex and diencephalon. We provide evidence that a major reason for these differences is the presence of Foxg1 in cortical cells, which reverses the cortical actions of Pax6 so that they occur in the same direction as in the diencephalon.

## MATERIAL AND METHODS

### EXPERIMENTAL MODEL

#### Mice colony maintenance and transgenic lines

To generate a conditional tamoxifen-inducible deletion of *Pax6* throughout the embryo, we combined lines carrying a *CAGGCre-ER^TM^* allele (Hayashi and McMahon, 2002), a green fluorescent protein (GFP) reporter allele (Sousa et al., 2009) and *Pax6^loxP^* alleles (Simpson et al., 2009). Pregnant mice were given 10mg of tamoxifen (Sigma) by oral gavage on embryonic day 9.5 (E9.5) to induce *Pax6^loxP^* deletion and embryos were collected on E11.5, E12.5 and E13.5. Embryos heterozygous for the *Pax6^loxP^* allele (*Pax6^fl/+^*;*CAGG^CreER^*) were used as controls since previous studies have shown no detectable defects in the forebrain of *Pax6^fl/+^* embryos (Simpson et al., 2009). Embryos carrying two copies of the floxed *Pax6* allele (*Pax6^fl/fl^*;*CAGG^CreER^*) were the experimental conditional knock-out (cKO) group. DTy54, a YAC transgene that expresses GFP in Pax6 expressing cells, was crossed into some animals to guide the diencephalic dissections (Tyas et al., 2006b).

To delete *Foxg1* in cortex, we combined lines carrying an *Emx1-CreER^TM^* allele (Kessaris et al., 2006), a GFP reporter allele (Sousa et al., 2009) and *Foxg1^loxP^* alleles (generously donated by Drs. Goichi Miyoshi and Gord Fishell; Miyoshi and Fishell, 2012). Pregnant females were given 10mg of tamoxifen by oral gavage at E9.5 and embryos were collected at E13.5 and E15.5. *Foxg1^fl/+^*;*Emx-CreER^TM^* embryos were considered controls and *Foxg1^fl/fl^*;*Emx-CreER^TM^* embryos were the experimental cKO group.

To simultaneously delete *Pax6* and *Foxg1* in the cortex, we combined lines carrying an *Emx1-CreER^TM^* allele (Kessaris et al., 2006), a GFP reporter allele (Sousa et al., 2009) and both *Pax6^loxP^* alleles (Simpson et al., 2009) and F*oxg1^loxP^* alleles (Miyoshi and Fishell, 2012). Pregnant females were given 10mg of tamoxifen by oral gavage at E9.5 and embryos were collected at E13.5 and E15.5.

The day the vaginal plug was detected was considered E0.5.

Animals were bred according to the guidelines of the UK Animals (Scientific Procedures) Act 1986 and all procedures were approved by Edinburgh University’s Animal Ethics Committee.

### METHOD DETAILS

#### Tissue processing for RNA-seq

We bisected embryonic brains (E13.5) along the midline and processed one half of the brain for immunohistochemistry to confirm Pax6 or Foxg1 loss in cKO samples. The other half was dissected further. For *Pax6*-cKO and corresponding control embryos, we dissected the thalamus (Th), prethalamus (PTh) and anterior half of the cerebral cortex (ACtx) (Figure 1-figure supplement 1) and extracted total RNA with an RNeasy Plus micro kit (Qiagen). We sequenced three biological replicates for ACtx and four biological replicates for Th and PTh. Each replicate consisted of samples pooled from three (ACtx and Th) or five (PTh) embryos of the same experimental group. We used embryos from 14 different litters, pooling control and experimental embryos from the same litter whenever possible. Poly-A mRNA was purified and TruSeq RNA-seq libraries were prepared and sequenced with Illumina HiSeq v4 (50 base paired-end reads for ACtx samples; 125 base paired-end reads for Th and PTh samples). For *Foxg1*-cKO and corresponding control embryos, we dissected only cortical tissue. Total RNA extraction and RNA-seq library preparation was performed as above (150 base paired-end reads). Five biological replicates were included, each consisting of two pooled cortices from littermate embryos.

We performed a post-processing quality control of samples through a series of principal component analysis (PCA) to see which samples satisfy the criteria of minimal within-group variance. Certain samples (1 PTh control, 1 Th control and 1 Th Pax6 cKO) did not satisfy the criteria since PCA resulted in them clustering out of their groups. We decided to remove these samples from further analysis in order to minimise unwanted technical variance.

#### Tissue processing for immunohistochemistry and *in situ* hybridization (ISH)

Embryos were dissected in cold phosphate buffered saline (PBS), their heads were fixed in 4% paraformaldehyde (PFA) overnight at 4°C, washed in PBS, cryoprotected in 30% sucrose and embedded in a mixture of 30% sucrose and OCT compound (50:50). Cryostat sections (5 or 10μm) were obtained and stored at −20°C until processed.

##### Immunohistochemistry

Cryo-sections were let to stabilize at room temperature for at least 2 hours and then washed three times in PBST (1X PBS with 0.1% Triton X-100, Sigma). To block endogenous peroxidase, sections were treated with 3% H_2_O_2_ for 10 minutes. After two PBS washes, antigen retrieval was performed by immersing the sections in Sodium Citrate buffer (10mM, pH6) heated at approximate 90°C using a microwave for 20 minutes. Once the solution cooled down sections were washed twice in PBST. After a 20 minutes pre-incubation in 20% Normal Goat Serum (Invitrogen), sections were incubated with the primary antibody overnight at 4°C. Biotin-coupled secondary antibodies were incubated for 1 hour at room temperature followed by a 30 minute incubation with Avidin-Biotin complex (ABC kit, Vector laboratories). Finally, diaminobenzidene (DAB, Vector Laboratories) reaction was used to obtain a brown precipitate and sections were mounted in DPX media (Sigma).

For immunofluorescence a cocktail of primary antibodies were incubated overnight at 4°C. Secondary antibodies were incubated at room temperature for one hour. For Ki67, Foxg1 and Tbr1 detection we used Streptavidin signal amplification (biotin-coupled secondary antibody followed by 30 minute incubation with Streptavidin Alexa Fluor™ 488, 546 or 647 conjugate; Thermo Fisher Scientific). Sections were counterstained with DAPI (Thermo Fisher Scientific) and mounted in ProLong Gold Antifade Mountant (Thermo Fisher Scientific).

Details of the antibodies used in this study can be found in Key Resources Table.

##### In situ hybridization

In vitro transcription of digoxigenin-labeled probes was done with DIG RNA-labeling kit (Sigma-Aldrich).

Cryo-sections were processed for ISH using standard protocols. Digoxigenin-labelled probes used were *Ccnd1* (kindly donated by Dr. Ugo Borello, INSERM, France), *Dlx2* (kindly donated by Dr. John L.R. Rubenstein, USCF, USA), *Neurog2* (kindly donated by Dr Thomas Theil, University of Edinburgh, UK), *Gbx2* (kindly donated by Dr. Alexandra L. Joyner, HHMI,USA), *Gsx2* (kindly donated by Dr. Kenneth Campbell, Cincinnati Children’s Hospital Medical Center, USA), *Dbx1* (kindly donated by Dr. Luis Puelles, University of Murcia, Spain), *Lef1* (kindly donated by Dr. J. Galcerán, University of Alicante, Spain), *Sfrp2* (kindly donated by Dr. Jeremy Nathans, JHU, USA), *Dkk3* (synthetized in the lab from cDNA using primers specified in Witte et al., 2009), *Foxg1* (kindly donated by Dr. Thomas Theil) and *Ascl1* (kindly donated by Dr. Francois Guillemot, Francis Crick Institute, UK).

Some slides were sequentially processed for fluorescent ISH (*Sfrp2*) followed by immunofluorescence (Pax6, Biolegend).

#### Genotyping of mutant lines

We dissected tissue from the tails of each embryo, extracted DNA and performed PCR amplification to detect the alleles of interest.

For the detection of the floxed Pax6 allele, PCR reaction was performed in a final volume of 25μl containing 1.5μl of extracted DNA, 0.5mM primer mix (Simpson et al. 2009, forward primer: 5’-AAA TGG GGG TGA AGT GTG AG-3’; reverse primer: 5’-TGC ATG TTG CCT GAA AGA AG-3’), 0.5 mM dNTPs mix, 1X PCR reaction buffer and 5U/μl Taq DNA Polymerase (Qiagen). PCR was performed with 35 cycles and a Tm of 59°C. The PCR product was subsequently run in a 2% agarose gel. Wild type allele results in a fragment of 156bp and floxed allele fragment was 195bp, therefore two bands indicated the heterozygous condition (used as controls) and one strong 195bp band identified the homozygous floxed allele condition (*Pax6* KOs).

For genotyping the floxed Foxg1 allele PCR reaction was performed in a final volume of 50μl containing 4μl of extracted DNA, 0.4mM primer mix (forward primer: 5’-TTGCTACATGCCTTGCCAG-3’; reverse primer: 5’-TCCAGCATCACCCAGGCGTC-3’), 0.2 mM dNTPs mix, 1X PCR reaction, 5% DMSO, and 5U/μl Taq DNA Poltymerase (Qiagen). PCR was performed with 34 cycles and a Tm of 58°C. The PCR product was subsequently run in a 2% agarose gel. Wild type allele results in a fragment of 190bp and floxed allele fragment was 230bp, therefore two bands indicated the heterozygous condition (controls) and one 230bp band identified the homozygous floxed allele condition (*Foxg1* KO).

#### Microscopy and imaging

ISH and DAB images were taken with a Leica DMNB microscope coupled to a Leica DFC480 camera. Fluorescence images were taken using a Leica DM5500B automated epifluorescence microscope connected to a DFC360FX camera. Images of embryo dissections were taken with a Leica MZFLIII fluorescence stereomicroscope. Image panels were created with Adobe Photoshop CS6.

#### Bromodeoxyuridine (BrdU) injections

Pregnant females, previously gavaged with tamoxifen at E9.5 to induce Pax6 deletion (see methods above), were intraperitoneally injected with a single dose of BrdU (10ug/ul, Thermo Fisher Scientific) at E10.5, E11.5 or E12.5 and embryos were collected 24 hours after the injection (E11.5, E12.5 or E13.5, respectively).

#### Quantitative Real Time PCR (qRT-PCR)

We extracted total RNA with RNeasy Plus micro kit (Qiagen) from Th, PTh and Ctx. cDNA was synthesized with a Superscript reverse transcriptase reaction (Thermo Fisher Scientific) and we performed qRT-PCR using a Quantitect SYBR Green PCR kit (Qiagen) and a DNA Engine Opticon Continuous Fluorescence Detector (MJ Research). We used the following primer pairs: Dlx2, 5’-CCAAAAGCAGCTACGACCT-3’ and 5’-GGCCAGATACTGGGTCTTCT-3’; Ngn2, 5’-CAAACTTTCCCTTCTTGATG-3’ and 5’-CATTCAACCCTTACAAAAGC-3’; Wnt8b, 5’-AACGTGGGCTTCGGAGAGGC-3’ and 5’-GCCCGCGCCGTGCAGGT-3’; Ccnd1, 5’-GAAGGGAAGAGAAGGGAGGA-3’ and 5’-GCGTACCCTGACACCAATCT-3’. We calculated and plotted the abundance of each transcript relative to GAPDH expression levels. For all experimental groups we used three biological replicates consisting on tissue dissected from embryos belonging to three independent litters. Controls and experimental embryos were from the same litter whenever possible. For each biological replicate we run three technical replicates by replicating the measurements three times and calculating the average.

### QUANTIFICATION AND STATISTICAL ANALYSIS

#### RNA-seq data analysis

##### Read alignment and counting

RNA-seq reads from each sample were mapped using STAR 2.4.0i (Dobin et al., 2013) to the mm10 mouse genome build downloaded from Ensembl77 (Aken et al., 2016) in October 2014. STAR was run with default options, allowing maximum multi-mapping to three sites. The number of reads mapped to each gene was counted using featureCounts v1.4.5-p1 (Liao et al., 2014) from the Subread package. Default options were used with a requirement that both reads needed to be properly mapped over exons to be counted, including reads which aligned over splice junctions.

##### Differential expression (DE) analysis

DE analysis was done in R (R Core Team, 2016) to assess the significance of differences in gene expression levels in cKO samples over control samples. We used two different R packages: DESeq2 1.8.1 (Love et al., 2014) and edgeR 3.1.12 (Robinson et al., 2009), both run with default parameters. DE was considered probable at FDR ≤ 0.05 for edgeR and an adjusted p-value ≤ 0.05 for DESeq2. Some genes were identified as differentially expressed by either DESeq2 alone or edgeR alone, while most of the genes identified as differentially expressed were identified by both (78.56% in ACtx, 73.96% in PTh and 70.60% in Th). After examining the expression levels of those genes by plotting counts per million (CPM) mapped reads in controls versus cKO, we decided to accept genes identified by either package as differentially expressed, attributing the difference in detection to the underlying methods (low-expression filtering, expression normalization and multiple testing correction) of each package. Accordingly, we included all genes identified by either DESeq2 or edgeR in our subsequent analysis.

##### Functional analysis

Lists of genes were analysed for Gene Ontology (GO) term enrichment using DAVID 6.8 Beta (Huang et al., 2009a, 2009b). Enrichment statistics were calculated for biological process terms from DAVID GO FAT database (GO_BP_FAT category).

##### Plots generated in R

Plots were generated with ggplot2 package (Wickham, 2016) and heatmaps were generated with pheatmap package. MA-plots were generated using the plotMA() function from DESeq2, modified so that it also includes the DE results from edgeR. PCA plots were generated using prcomp() function from base R.

#### Sample clustering methods

Expression values were transformed from raw read counts using variance stabilizing transformation described in DESeq2 and samples were hierarchically clustered using dist() and hclust() functions from base R, using parameters for Euclidean distance and Ward’s linkage method (Ward, 1963). To cluster samples from experiments in which *Pax6* or *Foxg1* were deleted, variance stabilizing transformation was followed by the application of the ComBat function from R package sva (Leek et al., 2016) to correct for batch effects that might influence the comparison of results from these two sets of experiments. These samples were clustered using the hcluster() function from amap R package (Lucas, 2014), with same distance and linkage methods as above. Within each tissue, log_2_-fold changes (LFCs) in gene expression between genotypes were calculated from average counts per million (CPM) mapped reads in cKO samples over average CPM mapped reads in control samples, using the following formula:

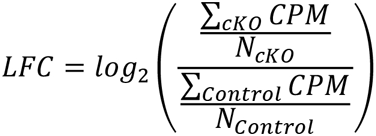

Genes were clustered hierarchically by Pearson’s correlation as a distance measure and Ward’s linkage method.

#### Image analysis and quantification

##### BrdU quantification and calculation of proliferation/differentiation indexes

We counted the total number of BrdU-labelled cells and classified them into three different categories according to their expression of Ki67 (proliferation category), Tuj1 (differentiation category) or none (G0 category). We calculated the fractions of proliferative cells, cells in G0 and differentiated cells by dividing the number of cells in each of the three categories by the total number of BrdU-labelled cells. Cell quantification was performed on 40x magnification coronal microphotographs using the cell counter plug in from Fiji (Image J) (Schindelin et al., 2012).

We quantified three biological replicates (three embryos from three different litters, n=3) for each tissue (Actx, Thal, Pthal), genotype (control, *Pax6* cKO) and age (E11, E12, E13), being controls and experimental embryos pairs from the same litter. For each embryo we analysed three different rostro-caudal sections in the case of cortical tissue and four rostro-caudal sections for diencephalic tissues. For each tissue and condition, we counted an average of 989 cells.

The data from all ages, regions and genotypes was statistically assessed to test the effects of Pax6 inactivation on proliferation and differentiation depending on age and tissue. Data was fitted to a generalized mixed linear model using the glmer() function from lme4 R package (Bates et al., 2015). Counts of cells in proliferation and counts of cells exiting the cell cycle were set as outcome variables, with genotype, age and tissue set as interacting fixed effects and litter and embryo set as nested random effects. Function argument ‘family’ was set to ‘binomial’ due to two possible outcomes of cell state. P-values of fixed effects and their interactions were obtained using the Anova() function from car package (Fox and Weisberg, 2011) with argument type = 3 to specify the usage of Type III Wald chisquare tests. Contrasts of interest were tested using lsmeans() function from lsmeans package (Lenth, 2016).

##### Quantification of proliferation

To quantify proliferation in the cortices of *Foxg1* cKOs and *Pax6 Foxg1* cKOs we selected an area of the cortex (indicated in Figure 9) and counted the total number of cells (DAPI-positive) and the number of proliferating cells (Ki67-positive). Proliferation fraction was calculated by dividing the number of proliferating cells by the total number of cells.

Cells were counted in 40x magnification coronal microphotographs using cell counter plug in from Fiji (Image J) (Schindelin et al., 2012). We quantified three independent embryos from three different litters for each genotype (n=3). For each embryo we counted three different rostro-caudal sections. Differences across the two genotypes were assessed by one tail paired t-test and significance was considered when p<0.05.

#### Statistical analysis /RT-qPCR details

T-tests (n=3) were performed in Microsoft Excel. Statistical details of all experiments are specified in the text or corresponding figure legend.

#### Analysis of the splicing variants Pax6 and Pax6 (5a) ratio

We used summarizeOverlaps() function from the GenomicAlignments R package (Lawrence et al., 2013) to count the number of reads aligning to genomic regions of Pax6 exons 5 and 5a. Read counts were divided by the number of bases of each exon (216b for exon 5, 42b for exon5a) to normalize for the exon length. Exon5/5a ratios for each sample were calculated as the normalized read counts in exon 5 over exon 5a. We then used aov() and TukeyHSD() functions from R stats package (base R) to test for significance in difference of exon5/5a rations between tissues and perform pairwise comparisons.

### RAW AND PROCESSED DATA AVAILABILITY

RNA-seq raw data of the Pax6 deletion experiment can be obtained from the European Nucleotide Archive (https://www.ebi.ac.uk/ena/data/view/PRJEB9747; Project 2015054, ENA accession numbers PRJEB9747, ERP010887).

RNA-seq raw data of the Foxg1 deletion experiment can be obtained from the European Nucleotide Archive (https://www.ebi.ac.uk/ena/data/view/PRJEB21349; Project 10900, ENA accession numbers PRJEB21349 and ERP023591).

To interactively explore the Pax6 RNA-seq dataset visit https://pricegroup.sbms.mvm.ed.ac.uk/Pax6_diencephalon/

---

**KEY RESOURCES TABLE.**
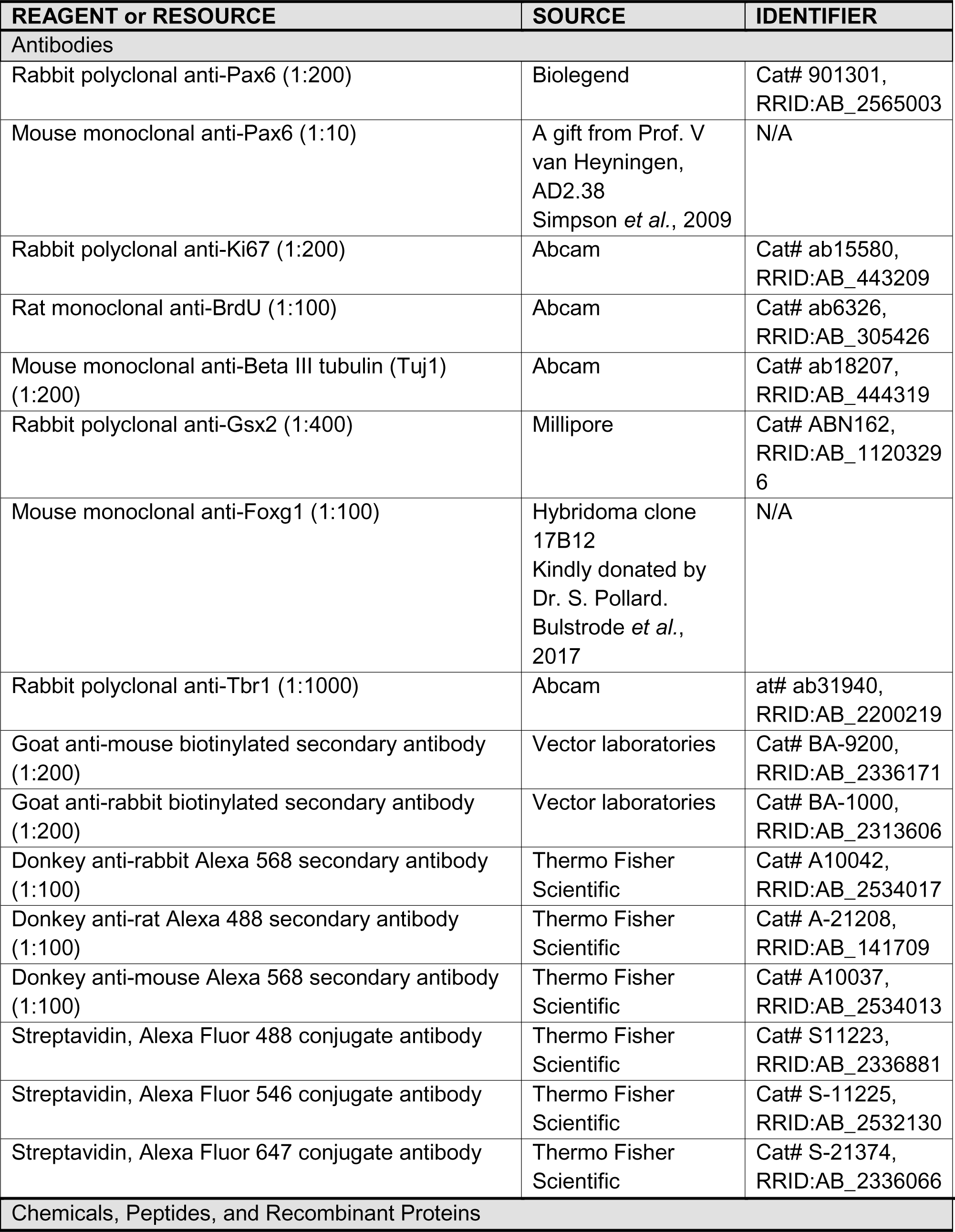
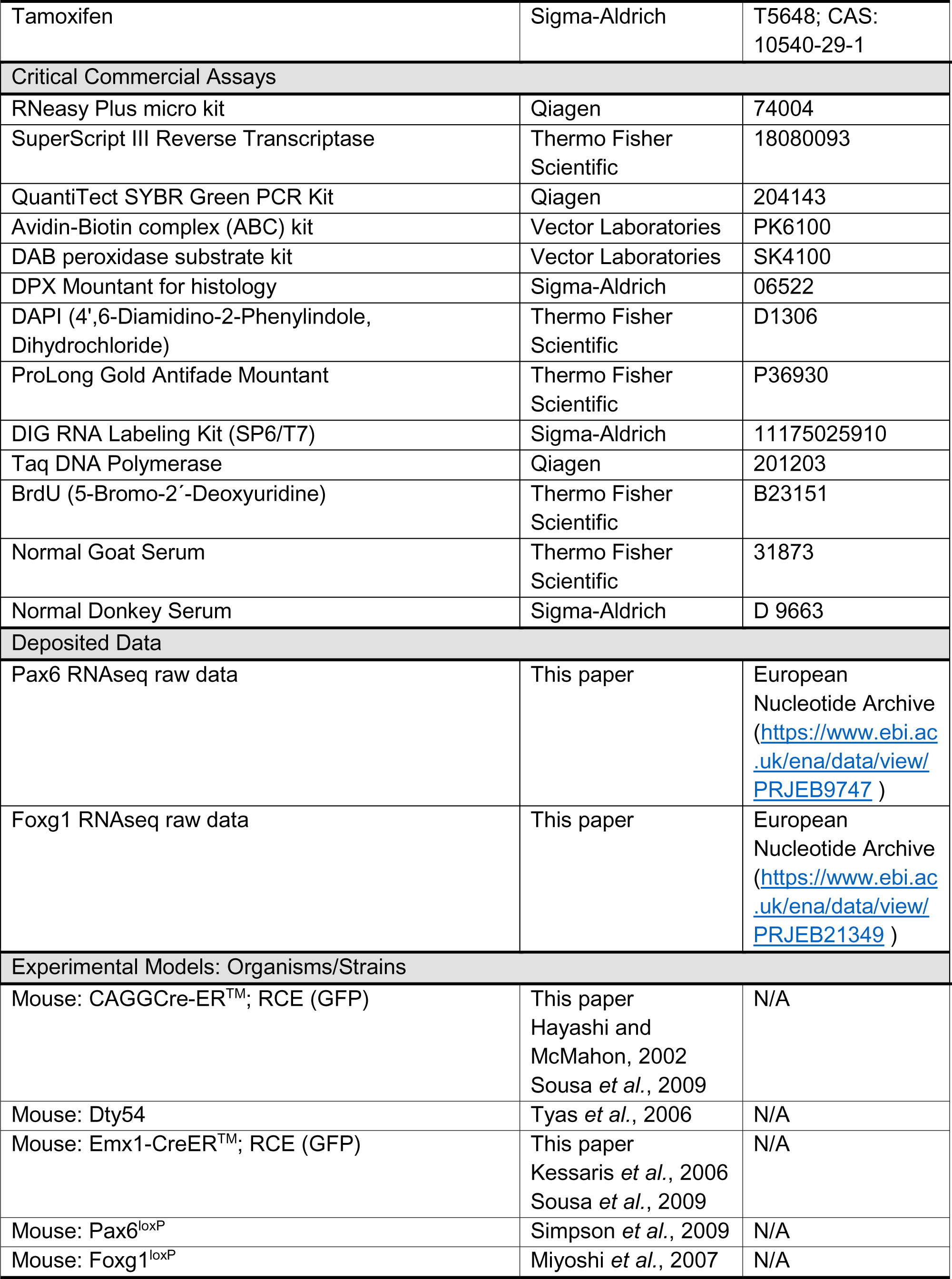
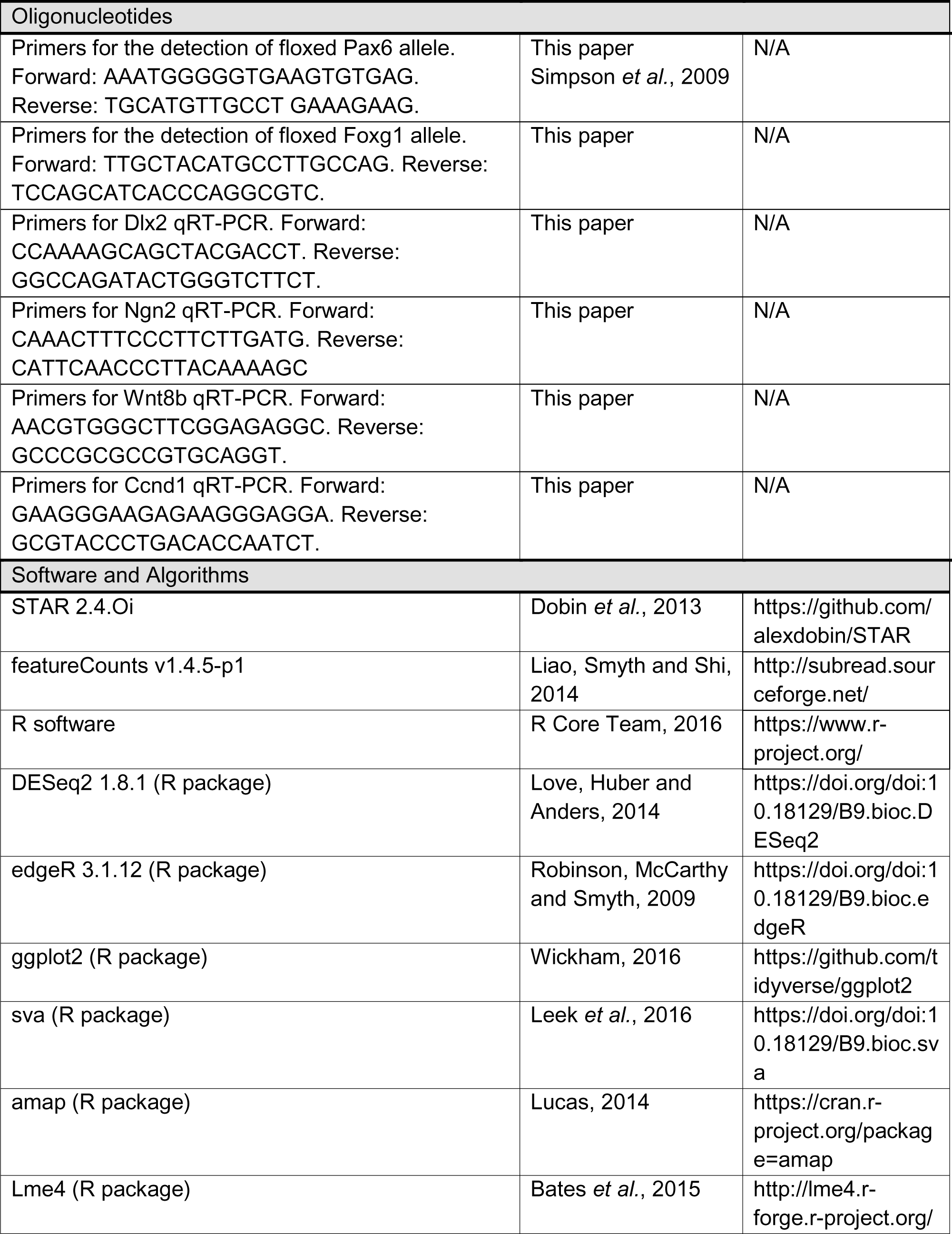
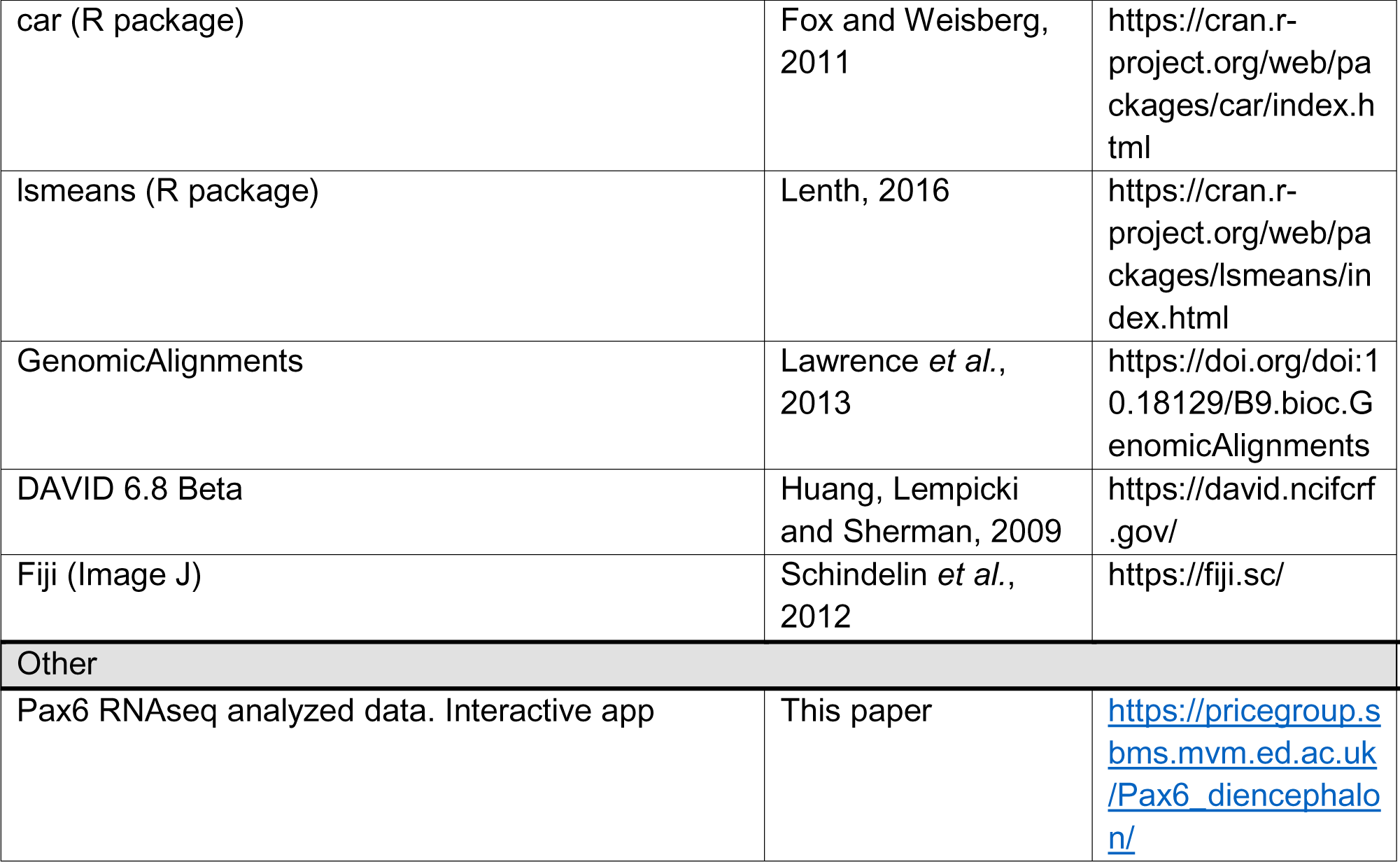

## Acknowledgements

An MRC UK Research Grant (N012291), a BBSRC UK Research Grant (N006542), a Royal Society UK Fellowship (NF151396) and a Marie Curie Fellowship EC (624441) funded the work; the *Foxg1^fl^* mice were a gift from Dr Goichi Miyoshi (Tokyo Women’s Medical University) and Dr Gordon Fishell (Harvard Medical School). RNAseq was performed at Edinburgh Genomics facilities (University of Edinburgh). We thank Chrysoula Giasafaki for her help in the Wnt study.

## Figure supplements legends

**Figure 1-figure supplement 1. Preparation of samples for RNA-seq.**

(A) Western blots showing Pax6 levels in two wild-type (WT), two *Pax6^+/-^* and a *Pax6^-/-^* E13.5 forebrain(s).

(B) GFP expression from the *DTy54* transgene in the E13.5 forebrain; Ctx, cortex; Di, diencephalon; Mes, mesencephalon.

(C,C’) The two cortices were teased apart and the brain was cut along the midline (broken line).

(D,D’) The anterior cortex (ACtx; high *Pax6*/GFP expression) was separated from posterior cortex (PCtx; broken line) in each hemi-brain. Th, thalamus; PTh, prethalamus.

(E,E’) Thalamus (Th) and prethalamus (PTh) were dissected (broken lines). Arrows indicate boundaries of these structures visible in bright-field. PT, pretectum.

**Figure 1-figure supplement 2. Expression patterns of genes showing the greatest inter-regional differential expression in control embryos. Related to Figure 1 and Supplementary File 1.**

(A,D,E,G,J,L,O,R-X,BB,CC,EE,GG,II) From Genepaint, http://www.genepaint.org/

(B,C,F,H,I,K,M,N,P,Q,Y-AA,DD,FF,HH,JJ) From Allen Brain Atlas http://developingmouse.brain-map.org/.

**Figure 2-figure supplement 1. Quality control of samples for RNA-seq.**

(A,B) Quantitative real-time PCR (qRT-PCR) to measure the levels of expression of *Dlx2* and *Neurog2* in E13.5 control thalamus (Th) and prethalamus (PTh) relative to *GAPDH*. Values are means ± sem; n=3 animals in all cases (p<0.05 Student’s t-test).

(C-H) In situ hybridizations on control forebrains at E13.5 and data from control and *CAG^CreER^ Pax6* cKOs on counts per million reads (CPM) extracted from RNA-seq experiments for three thalamic and three prethalamic markers. Red arrows indicate low values for markers of each region in the other region.

This figure is linked to Figure 1 and material and methods section.

**Figure 2-figure supplement 2. Principal component analysis on RNA-seq data.**

(A-C) Analysis of data from 3 samples from each of control and *CAG^CreER^ Pax6* cKO anterior cortex, control prethalamus and control and *CAG^CreER^ Pax6* cKO thalamus and 4 samples from *CAG^CreER^ Pax6* cKO prethalamus.

**Figure 4-figure supplement 1. Effects of Pax6 deletion on proportions of different cell types in forebrain regions with age in control and *CAG^CreER^ Pax6* cKOs.**

Counts in ACtx were in lateral (L) and medial (M) regions. Proliferating cells were BrdU+, Ki67+, Tuj1-; differentiating cells were BrdU+, Ki67-, Tuj1+; intermediate cells were BrdU+, Ki67-, Tuj1-.

Three biological replicates were included (three embryos from three different litters, n=3) for each tissue (Actx, Thal, Pthal), genotype (control, *Pax6* cKO) and age (E11, E12, E13), being controls and experimental embryos pairs from the same litter. Bars indicate SEMs.

**Figure 7-figure supplement 1. Effects of Foxg1 deletion from the cortex.**

(A) Principal component analysis of RNA-seq data from 5 control and 5 *Foxg1* cKO samples.

(B) MA plots of log_2_ fold changes in the expression of each gene against its average expression level; red dots indicate statistically significant changes between genotypes (adjusted p values <0.05).

(C) Numbers of significantly (adjusted p<0.05) upregulated and downregulated genes in *Foxg1* cKO cortex.

(D) Expression of Foxg1 and Pax6 in *Foxg1* cKO. Panels show merged, Pax6 and Foxg1 staining. Scale bar: 0.25mm.

(E) Counts per base read coverage of exons 4, 5a and 5 of *Pax6* from each control and *Foxg1* cKO sample.

(F) Ratios between the average coverage per base in exons 5 and 5a for each sample superimposed on box and whisker plots. There was no significant difference between genotypes.

**Figure 7-figure supplement 2. Clustering of RNA-seq data on genes annotated by the Neuron Differentiation and Wnt signalling pathway GO terms.**

(A,C) Heatmaps of hierarchical clustering of RNA-seq data from samples of control cortex, control ACtx, control PTh, control Th and *Emx1^CreER^ Foxg1* cKO cortex.

(B,D) Principal component (PC) analysis on the same RNA-seq data as in A,C).

## Supplementary Files

**Supplementary File 1.** Lists of genes showing significant enrichment in each control tissue over the other two (adjusted p<0.05). Linked to Figure 1.

**Supplementary File 2.** Lists of genes showing significant enrichment in each *CAG^CreER^ Pax6* cKOs tissue over the other two Pax6 cKOs (adjusted p<0.05). Linked to Figure 1.

**Supplementary File 3.** List of genes regulated in opposite directions in anterior cortex versus thalamus, anterior cortex versus prethalamus or thalamus versus prethalamus Pax6 cKOs tissues. Linked to Figure 1.

**Supplementary File 4.** Genes showing significant differential expression between controls and Pax6 cKOs (adjusted p<0.05) for each tissue analysed (anterior cortex, thalamus and prethalamus). Linked to Figure 2.

**Supplementary File 5.** Genes showing significant upregulation or downregulation in all three tissues (anterior cortex, thalamus and prethalamus). Linked to Figure 2.

**Supplementary File 6.** Full list of functional terms associated with each cluster shown in Figure 3A. Linked to Figure 3.

**Supplementary File 7.** Genes showing significant differential expression in *Foxg1* cKOs (*Emx1^CreER^*;*Foxg1^fl/fl^*). Adjusted p<0.05. Linked to Figure 7.

**Supplementary File 8.** Functional terms showing enrichment in upregulated and downregulated genes in *Foxg1* cKOs. Adjusted p<0.05. Linked to Figure 7.

**Supplementary File 9.** Significant deregulated genes in *Foxg1* cKO (adjusted p<0.05) annotated by the GO term “Cell Cycle” used for the hierarchical clustering showed in Figure 7. Linked to Figure 7.

**Supplementary File 10.** Genes showing significant deregulation in both *Foxg1* cKOs and *Pax6* cKOs cortices. Linked to Figure 8.

